# A web-based histology atlas for the freshwater sentinel species *Daphnia magna*

**DOI:** 10.1101/2022.03.09.483544

**Authors:** Mee S. Ngu, Daniel J. Vanselow, Carolyn R. Zaino, Alex Y. Lin, Jean E. Copper, Margaret J. Beaton, Luisa Orsini, John K. Colbourne, Keith C. Cheng, Khai C. Ang

**Affiliations:** Department of Pathology, Pennsylvania State University College of Medicine, Pennsylvania, USA; Jake Gittlen Laboratories for Cancer Research, Pennsylvania State University College of Medicine, Pennsylvania, USA; Department of Biology, Mount Allison University, Sackville, Canada; Centre for Environmental Research and Justice, The University of Birmingham, Birmingham, UK; Institute for Computational and Data Sciences, Pennsylvania State University, State College, Pennsylvania, USA; Molecular and Precision Medicine Program, Pennsylvania State University College of Medicine, Hershey, Pennsylvania, USA

**Keywords:** *Daphnia magna*, sentinel, microanatomy, atlas, phenotypes, sexual dimorphism, histopathology, toxicology

## Abstract

*Daphnia* are keystone species of freshwater habitats used as model organisms in ecology and evolutionary biology. Their small size, wide geographic distribution, and sensitivity to chemicals make them useful as environmental sentinels in regulatory toxicology and chemical risk assessment. Biomolecular (-omic) assessments of responses to chemical toxicity, which reveal detailed molecular signatures, become more powerful when correlated with other phenotypic outcomes (such as behavioral, physiological, or histopathological) for comparative validation and regulatory relevance. However, the lack of histopathology or tissue phenotype characterization of this species presently limits our ability to access cellular mechanisms of toxicity. Here, we address the central concept that interpreting aberrant tissue phenotypes requires a basic understanding of species normal microanatomy. We introduce the female and male *<u>Da</u>phnia* <u>H</u>istology <u>R</u>eference <u>A</u>tlas (DaHRA) for the baseline knowledge of *Daphnia magna* microanatomy. Additionally, we also included developmental stages of female *Daphnia* in this current atlas. This interactive web-based resource of adult *Daphnia* features overlaid vectorized demarcation of anatomical structures whose labels comply with an anatomical ontology created for this atlas. We demonstrate the potential utility of DaHRA for toxicological investigations by presenting aberrant phenotypes of acetaminophen-exposed *D. magna*. We envision DaHRA to facilitate the effort of integrating molecular and phenotypic data from the scientific community as we seek to understand how genes, chemicals, and environment interactions determine organismal phenotype.

## 1. Introduction

Environmental pollution is the leading cause of premature morbidity and mortality globally (Fuller et al., 2022; Landrigan et al., 2018; Naidu et al., 2021). Habitat loss, climate change, and pollution also impact biodiversity, with more than 60% of ecosystems services being diminished in the last two decades (Cardinale et al., 2012). New approach methodologies (Stucki et al., 2022) for assessing chemical toxicity are developed to improve regulatory outcomes by replacing outdated, data-poor methods dependent upon apical endpoints (such as death or reproductive failure) with data-rich molecular data (e.g. transcriptomic and metabolomic) (Harrill et al., 2021; Hines et al., 2010; Palmer et al., 2020). These data are robust at measuring biomolecular activity that are potentially indicative of chemical modes of action. However, these data lack spatial resolution, context within the whole organism, or correlation with associated abnormal tissue phenotypes that can be highly informative with regard to potential human toxicity (European Chemicals Agency, 2020). Since understanding abnormal tissue phenotypes requires knowledge of normal microanatomy (microscopic anatomy or histology), the primary objective of this paper is to provide a resource for visualizing and interpreting tissue phenotypes for the model species *D. magna* – an organism used globally to set regulatory limits on potentially hazardous chemical substances in the environment (United States Environmental Protection Agency, 1996, 2002; Organisation for Economic Co-operation and Development, 2004, 2018), and one of five models being used to uncover evolutionarily conserved toxicity pathways (“The Precision Toxicology initiative,” 2023).

The water flea *Daphnia* is a keystone branchiopod crustacean (order *Cladocera*) in freshwater lotic ecosystems worldwide and an established model in ecology, evolutionary biology, and ecotoxicology (Ebert, 2022; Miner et al., 2012; Stollewerk, 2010). They are responsive to environmental change and adapt via evolutionary mechanisms and plasticity (Cuenca-Cambronero et al., 2021; Stoks et al., 2016; Walsh et al., 2018). Relevant to ecotoxicity testing is their short generation time that enables the experimental manipulation of large populations and a parthenogenetic life cycle that allows the rearing of populations of identical clones (Hebert and Ward, 1972). The latter property has the unique advantage of facilitating the concurrent study of molecular and phenotypic responses to multiple environmental insults, including chemical pollutants (Abdullahi et al., 2022; Cuenca Cambronero et al., 2018). *Daphnia magna* is a model species for ecotoxicogenomics (Kim et al., 2015; Shaw et al., 2008). Recently, its hologenome (Chaturvedi et al., 2023), reference genome (Byeon et al., 2022; Lee et al., 2019) and transcriptome (Campos et al., 2018; Jankowski et al., 2022; Orsini et al., 2016) have been published, elevating this species to the ranks of other biomedical model species for ecological genomics. The full potential of this species is best realized when correlations can be established between molecular, and tissue- and cell-specific phenotypes.

Histopathology, the microscopic examination of diseased tissues, enables the identification of targets of toxicity and diseases, bridging phenotypes and biomolecular perturbations induced by environmental insults (Majno and Joris, 2004; Wester and Canton, 1991). Histopathology-based toxicological studies in fish (Huang et al., 2021; Manjunatha et al., 2022; Ramírez-Duarte et al., 2008) and bivalves (Fraga et al., 2022; Joshy et al., 2022) have been useful for water quality monitoring and assessment. The application of histopathology to millimeter-size sentinel species used in ecotoxicology would enable the analysis of tissue-specific toxicity phenotypes in the context of the whole animal. However, identification of affected cell and tissue types requires prior knowledge of normal microanatomy; the use of web-based atlases maximizes accessibility across research and educational communities (Copper et al., 2018; Graham et al., 2015; van der Ven et al., 2003).

Here we present the first curated web-based female and male <u>D</u>aphnia <u>H</u>istology <u>R</u>eference <u>A</u>tlas (DaHRA; RRID:SCR_024913), further broadening the discovery capacity of this sentinel species. We have optimized methods for *D. magna* histology and created a collection of digitized histological images for adult female and male *D. magna* in three anatomical planes to illustrate sexual dimorphism associated with environmentally induced phenotypic plasticity. We also present a subset of developmental stages showcasing some representative developmental events. As proof-of-concept, we also present histological alterations in *D. magna* caused by exposure to toxic levels of a common pharmaceutical painkiller, acetaminophen, to demonstrate how our platform can facilitate whole-organism visualization and comparison for an experiment. This resource is made open-access and interactive, allowing smooth magnification with a dynamic scale bar. Anatomical structures are highlighted and labeled in compliance with an anatomical ontology we generated for the atlas, providing researchers and chemical risk managers with an unprecedented tool to navigate the microanatomy of *D. magna*. This atlas has the potential to support both tissue-specific and whole-organism phenotyping, informing (eco)toxicology, genetic and phenomic studies.

## 2. Material and methods

### 2.1 *Daphnia magna* culturing

A commercial clone of *D. magna* was purchased from Carolina Biological (NC, USA) and raised in “Aachener Daphnien-Medium” or ADaM at room temperature (20°C ± 1°C) under a 16-hours light/8-hours (hrs) dark photoperiod. *D. magna* cultures were fed three times weekly with 3.0 × 10^7^ cells/ml of green microalgae (*Raphidocelis subcapitata)* and once a week with 0.1 mg/mL of dissolved bakers’ yeast. The animal density was maintained at about 20 neonates, 10 juveniles and 6 to 8 reproducing adults per liter to prevent overcrowding. Under these conditions, animals reached maturity at 6 to 8 days post-birth and reproduced parthenogenetically every 3 days after sexual maturation with an average of 15 neonates per brood from the second brood onwards. Production of males was induced by overcrowding (>10 reproducing adults per liter) and shorter photoperiod (8 hrs) (Zhang and Baer, 2000).

### 2.2 Chemical Exposure

In order to have pronounced abnormal tissue phenotypes as a proof-of-concept to demonstrate the utilization of this atlas, we used a wide range of Lethal Concentration (LC) 50 values documented in the literature for acetaminophen (de Oliveira et al., 2016; Du et al., 2016). Reproducing female *D. magna* (approximately 10 days old and carrying the second parthenogenetic brood 2 hrs post-ovulation) were exposed to 5 concentrations of acetaminophen (5, 15, 25, 35, 50 ug/mL). Gravid *D. magna* were used for this exposure to evaluate the toxic effects of acetaminophen on both adults and developing embryos. The exposures lasted for 72 hrs and were conducted with two adult females in 200 ml medium. The medium was replenished, and the animals fed daily. After 72 hrs exposure, each surviving animal was prepared for histological observations as described in the following.

### 2.3 Histological Processing

#### 2.3.1 Fixation and decalcification

Exposed and control clones of *D. magna* were fixed with 20X Bouin’s solution (Newcomer Supply, WI) and incubated for 48 hrs at room temperature (about 21°C) on a low-speed orbital shaker (Corning LSE) set to 55 revolutions per minute (RPM). The fixation is done to preserve tissues from decay due to autolysis or putrefaction. After the fixation step, samples were washed twice with 1X phosphate-buffered saline (PBS) for 10 min. This washing step was followed by decalcification in 20X sample volume of pre-chilled 6% formic acid (Sigma-Aldrich, MO) for 24 hrs on the orbital shaker set to 55 RPM. Samples were then rinsed in 70% ethanol for one minute and immersed in fresh 70% ethanol for 30 min before agarose embedding. We tested fixation using 4% Paraformaldehyde (PFA) in 0.1M phosphate buffer (pH 7.4) (Bioenno LifeSciences, CA) and 10% Buffered Formalin Phosphate (NBF; Fisher Scientific, ON) with different fixation times and temperatures (See Table S1, Supplementary Material). The *D. magna* samples fixed using PFA and NBF (n=23) showed “ballooning”, a severe fixation artifact causing the carapace to ‘puff-up’ (See Figure S1, Supplementary Material). It was concluded after comparison of histological sections generated using these fixatives that Bouin’s solution is the best fixative for *D. magna* and was used to fix all samples used in this atlas (See Figure S2, Supplementary Material).

#### 2.3.2 Agarose embedding

Visualization of histological sections in each of the three standard anatomical planes (coronal, sagittal, and transverse) is critical for understanding organismal anatomy. Therefore, the ability to generate consistent sections in each of these planes is essential. Agarose embedding using a mold or an array facilitates consistent positioning and orientation of millimeter-size samples for sectioning (Copper et al., 2018; Santana et al., 2023). A mold was designed (See Figure S3, Supplementary Material) and 3D-printed for casting an agarose block with triangular wells that could hold up to 18 adult *D. magna* for concurrent tissue processing and sectioning (See Figure S3, Supplementary Material). To create an agarose block, laboratory labeling tape (VWR) was wrapped tightly around the mold. Then, 2.5 mL of 1 % agarose (Sigma-Aldrich, MO) at 55 °C was pipetted onto the mold and allowed to solidify at room temperature. The agarose block was removed gently from the mold. Each fixed *D. magna* sample was pipetted with a small volume of ethanol and transferred into the well of the agarose block using a single-use plastic transfer pipette. Samples designated for the sagittal plane sectioning were laid on their sides with a swimming antenna in the wells and all rostra facing the same direction (see Figure S4 for *Daphnia* anatomy and File S1 for *Daphnia* anatomy glossary, Supplementary Material). Samples designated for coronal and transverse orientation were laid on their back in the wells. Once all samples were positioned in individual wells, excess ethanol was carefully dried off using lint-free Kimwipes without touching the samples. Each sample was first topped off with one drop of molten 1 % agarose (about 50 °C) without disturbing the sample, followed by a thin layer of 1% agarose to completely cover the sample. After the agarose layer solidified (about 5 min at room temperature), the block was trimmed as needed, placed into a tissue cassette, and stored in 70 % ethanol for tissue processing.

#### 2.3.3 Tissue processing, sectioning, and staining

All samples were dehydrated in graded ethanol and infiltrated with Formula R paraffin (Leica Biosystems #3801450) in RMC Model 1530 automated closed reagent type tissue processor (See Table S2, Supplementary Material). Following this step, they were serially sectioned at 5 μm on a Leica RM2255 automated rotary microtome. Sections were then stained with Harris’ hematoxylin and eosin in an auto-stainer (Sakura Tissue Tek DRS 2000, IMEB, CA) following a protocol adapted from Copper et al. (2018) where the duration of hematoxylin staining was extended from 3 to 7 min to achieve better contrast for *Daphnia* samples (See Table S3, Supplementary Material). Cover glasses No. 1 (Platinum Line) were used for cover-slipping.

### 2.4 Histology slide digitization

All slides were screened using an Olympus BX41 microscope and 10X and 20X objective lenses. Those selected for the atlas were scanned at 40X using an Aperio AT2 slide scanner (Leica Biosystems, IL) and images were saved in TIFF format. 40X scanning was performed using 20X objective lens (0.075 n.a. Plan Apo) with 2X optical magnification changer, yielding a digital resolution of 0.25-micron per pixel. The images of *D. magna* samples included in the atlas were cropped using Aperio ImageScope (version 12.4.3.5008). Three channels (Red, Green, Blue) of these digital slides were stacked using Fiji (Schindelin et al., 2012) or ImageJ (Schneider et al., 2012). Then, image processing was performed in Adobe Photoshop (version 22.1.1) where images were rotated and set to have the same canvas size; the image background was removed using “Remove Background”; the “Exposure” was adjusted to fall between 0.1 to 0.25 and the same value was used for each set of images; and “Levels” were adjusted using preset “Midtone Darker”. Each set of digital slides was then pyramidally tiled for the web-based viewer.

### 2.5 Digital labeling of anatomical structures

The anatomical ontology, consisting of a list of anatomical terms organized by groups (organ systems) and subgroups (tissues and cell types), was created for the atlas (See Table S4, Supplementary Material). We cross-referenced the extensive work of Fryer (Fryer, 1991) with other published literature (Agar, 1950; Auld et al., 2010; Bednarska, 2006; Benzie, 2005; Binder, 1931; Christensen et al., 2018; Consi et al., 1987; Ebert, 2005; Edwards, 1980; Goldmann et al., 1999; Halcrow, 1976; Hiruta and Tochinai, 2014; Kikuchi, 1983; Kress et al., 2016; McCoole et al., 2011; Metschnikoff, 1884; Quaglia et al., 1976; Rossi, 1980; Schultz and Kennedy, 1976; Smirnov, 2013; Stein et al., 1966; Steinsland, 1982; Weiss et al., 2012; Wuerz et al., 2017; Zaffagnini and Zeni, 1987, 1986; Zeni and Franchini, 1990) and decided on the commonly used *Daphnia* anatomical terms. Annotation and labels for each anatomical structure presented on the atlas were created using Adobe Illustrator (version 25.1). One image at a time, each anatomical structure was annotated by outlining the structure using the “Curvature” and assigned a color corresponding with the anatomical ontology. Annotation and labels of each structure were saved under “Layers”. After completion of the labeling of all anatomical structures on a given image, the annotations were exported in single scalable vector graphic (SVG) file format to be used as input for the web-based viewer.

### 2.6 Building the web-based digital slide visualization platform

To improve accessibility and usability, we developed an open-access, web-based digital slide viewing platform based on the open-access project OpenSeadragon (https://openseadragon.github.io/). This interface removes the need to download full-resolution images. The viewer combines SVG files and digital scans into a seamless experience to provide user-friendly access to high-resolution data. The atlas’ code was written in client-side JavaScript, HTML, and CSS. Pyramidally tiled images were parsed and visualized with OpenSeadragon. When the user opens an image, the viewer opens the corresponding SVG file containing all the anatomical labels and their corresponding annotations. The viewer parses all labels from the SVG file, plots the corresponding regions, and updates the ontology to note which regions are available on a particular image.

## 3. Results and discussion

### 3.1 *<u>Da</u>phnia* <u>H</u>istology <u>R</u>eference <u>A</u>tlas (DaHRA) presenting *D. magna* microanatomy

#### 3.1.1 Interactive viewer

We developed the web-based atlas, DaHRA (http://daphnia.io/anatomy/), to be a user-friendly interface to access a collection of digitized histological sections of wildtype female and male *D. magna* in each of three standard anatomical planes (Figure 1A). DaHRA’s interface allows users to visualize digital scans of whole-organism sections up to 40X objective magnification (0.25- micron per pixel resolution), providing sufficient resolution to recognize virtually all cell types with broader organismal context. Compared to histology atlases of other model organisms (for example, zebrafish (Copper et al., 2018) and mouse embryos (Armit et al., 2017)), DaHRA offers interactive visualization of normal microanatomy using overlaid vectorized demarcation of anatomical structures whose labels comply with an anatomical ontology created for this atlas. The anatomical ontology consisting anatomical structures can be found on the left side of the viewer, with the anatomical terms arranged alphabetically within groups (Figure 1B). Annotations of the anatomical structures are presented as color overlays and indicated by check marks next to the anatomical terms. Unchecking the box hides the color overlays. Anatomical terms with underlined labels indicate nested substructures (for example, “microvilli” under “epithelial cell”, both under “midgut”). Hovering over an anatomical term in the ontology dynamically highlights the corresponding structure or structure groups in the viewer, temporarily hiding other checked structures.

**Figure 1.**
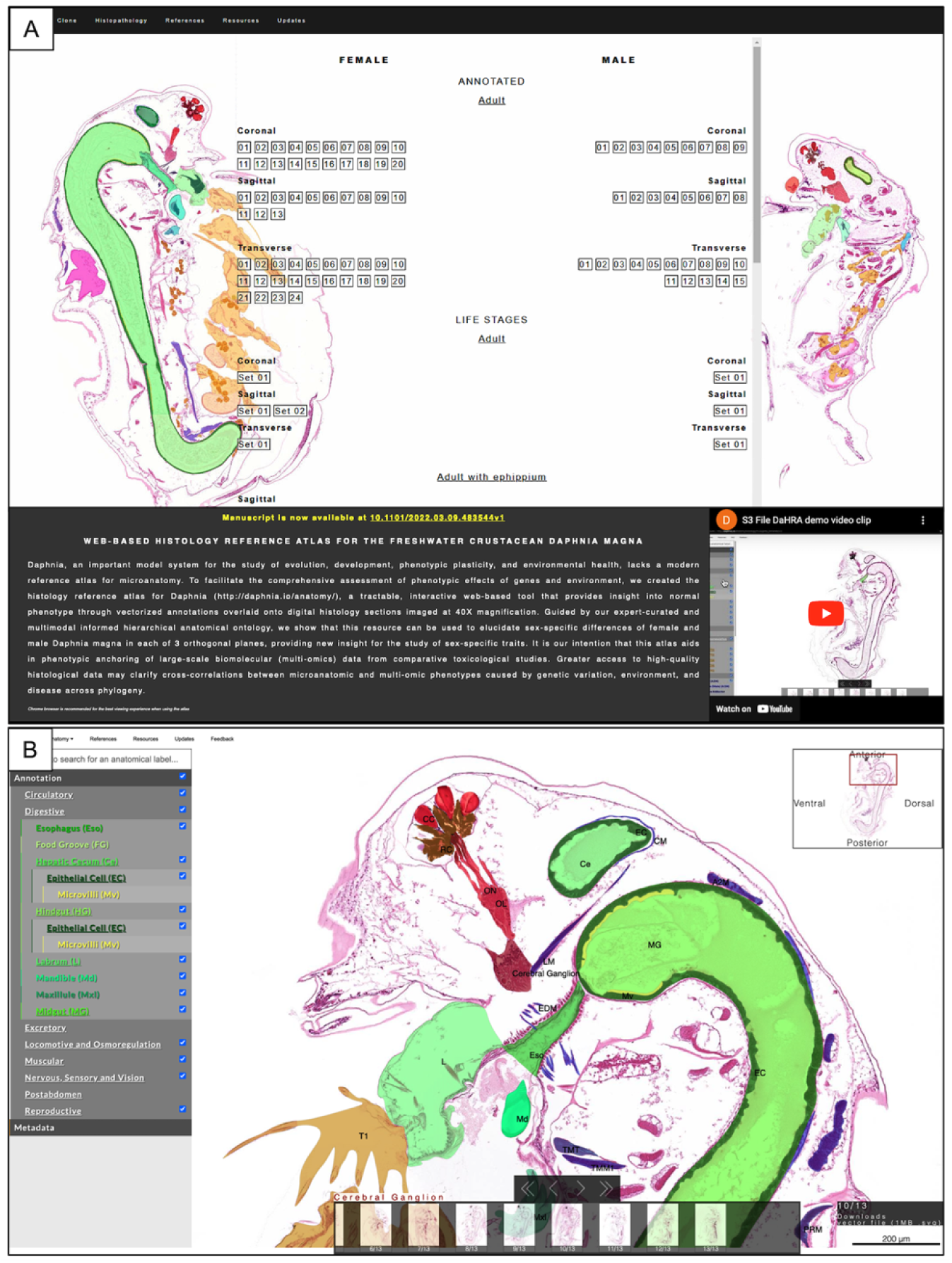
Interactive web-based viewer for DaHRA. **(A)** The landing page hosts annotated images of female and male commercial clone of D. magna with an instruction video describing the features of the atlas. Unannotated images of embryos at different stages, juvenile, and adults are also categorized under “Life stages”. **(B)** Interactive viewer displaying the expandable list of anatomical structures on the left; the checked boxes indicate the structures labeled in the image. The anatomical terms on the image are shown as acronyms to minimize obscuring of structures; hovering the mouse cursor over an acronym or its corresponding region will show the full term. Unchecking a box will hide the color overlay and annotation corresponding to the box.

In order to make the atlas as a central resource for *Daphnia* community, we also created “Reference” tab which lists published literature related to *Daphnia*’s specific organs or cell-types that were used to annotated this atlas, and “Resources” tab which contains a file annotating *Daphnia* gross normal anatomy, *Daphnia*-specific anatomical definitions, anatomy ontology curated for the atlas, and histology protocols optimized for *Daphnia* samples, including a casting mold stereolithography file . To facilitate collaboration, updates, and validation, a “Feedback” tab is provided for users to leave comments and suggestions. A video demonstrating the features of the atlas is also available on the atlas histology landing page.

#### 3.1.2 *D. magna* male and female microanatomy

DaHRA presents the first microanatomical representation of female and male *D. magna* from the same clone. A hundred samples were sectioned for protocol optimization, and one sample per orientation of each sex was selected and annotated for the atlas. Scanned images of 10 *D*. *magna* are presented in the histology atlas. Organs and cell types included in the anatomical ontology are briefly described here with representative images from the three anatomical planes of the female (Figures 2A-C) and male *D. magna* (Figures 2D-F). The terminology used for the DaHRA anatomical ontology (a list of terms organized by groups and subgroups) was cross-referenced with published literature (Agar, 1950; Auld et al., 2010; Bednarska, 2006; Benzie, 2005; Binder, 1931; Christensen et al., 2018; Consi et al., 1987; Ebert, 2005; Edwards, 1980; Fryer, 1991; Goldmann et al., 1999; Halcrow, 1976; Hiruta and Tochinai, 2014; Kikuchi, 1983; Kress et al., 2016; McCoole et al., 2011; Metschnikoff, 1884; Quaglia et al., 1976; Rossi, 1980; Schultz and Kennedy, 1976; Smirnov, 2013; Stein et al., 1966; Steinsland, 1982; Weiss et al., 2012; Wuerz et al., 2017; Zaffagnini and Zeni, 1987, 1986; Zeni and Franchini, 1990) for uniformity. We identified 50 anatomical structures and categorized them into 8 groups (circulatory, digestive, excretory, locomotive and respiration, muscular, nervous, sensory and vision, postabdomen, and reproductive), and can be expanded if/ when more structures are identified.

**Figure 2.**
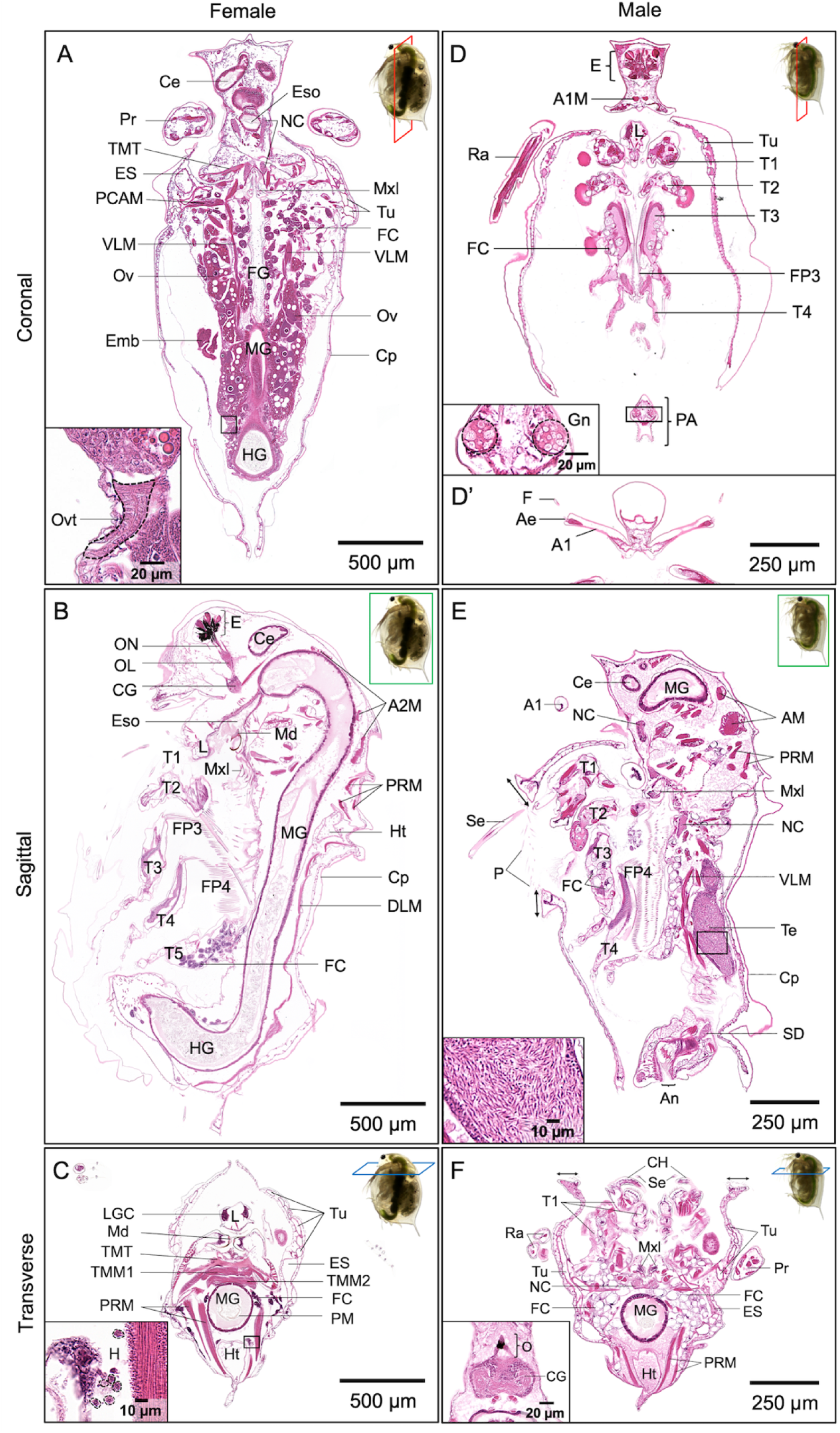
Representative microanatomical structures of female (A-C) and male (D-F’) *D. magna* in three orthogonal planes. The coronal plane (panels A and D) displays most of the structures in pairs. Inset of panel A shows the oviduct (Ovt; dotted circle) and inset of panel D shows gonopores (Gn) in the male. The histology section for Panel D’ is slightly ventral to that of panel D and displays prominent and elongated antennules (A1) with flagella (F) at the tips. The mid-sagittal plane of the female (panel B) includes connections between the compound eye (E) and the optic lobe (OL) and cerebral ganglia (CG) by optic nerves (ON). The labrum (L), maxillules (Mxl), and mandibles (Md) are anterior to the esophagus (Eso) that opens into the midgut (MG) and is followed by the hindgut (HG). This section also cuts through the five thoracic limbs (T1-5) and filter plates (FP3, FP4). The sagittal plane of the male (panel E) shows the elongated seta (Se) on the first thoracic limb, pubescence (P) at the wider ventral opening of the carapace, thickening of carapace at the ventral opening (arrows), one of the testes (Te), and a small portion of sperm duct (SD). Inset of panel E showing the spermatozoa in the testis. The transverse plane of the female (panel C) shows the asymmetrical paired mandibles (Md) with the transverse mandibular tendons (TMT), transverse mandibular muscles (TMM1), transverse muscles of mandibles (TMM2), and the posterior rotator muscles of mandibles (PRM). Inset of panel C displays several hemocytes (H) outlined by dotted circles. The transverse plane of male (panel F) displays the paired copulatory hooks (CH) on the first thoracic limbs (T1) and the thickening of the carapace (arrows) at the ventral opening. This also shows the abundance of fat cells (FC) which are quite different from those in the female. The histology section for Panel F’, is slightly anterior to that of panel F and shows the pigmented ocellus (O) is connected to the cerebral ganglia (CG). A1M, antennule muscle; A2M, antennal muscle; Ae, aethetasc; An, anus; Ce, hepatic cecum; Cp, carapace; DLM, dorsal longitudinal muscle; ES, end sac of the maxillary gland; FG, food groove; Ht, heart; LGC, labral gland cell; NC, nerve chord; PA, postabdomen; PCAM, posterior carapace adductor muscle; PM, peritrophic membrane; Ra, ramus of swimming antenna; Tu, tubule of the maxillary gland; VLM, ventral longitudinal muscle. Atlas links to each panel can be found in File S2, Supplementary Material.

We first summarize the sexually dimorphic traits of *D. magna* and follow with brief descriptions of the normal anatomy and microanatomy. Adult males have a smaller body size and much longer antennules than females, that bear a single long flagellum on the tip (Benzie, 2005)(Figure 2D’). Male antennules contain muscle tissue (Figure 2D) that is absent in females. The first thoracic limbs of the males are equipped with elongated setae (Figure 2E) and chitinized copulatory hooks (Figure 2F) that are used for clasping females during copulation. The male postabdomen has gonopores (Figure 2D inset) that are involved in transferring mature spermatozoa from the testes to the female in the region of the oviduct during copulation. Besides having a wider frontal opening, pubescence and thickened angular margins are also observed at the ventral margin of the carapace in males (indicated by arrows in Figures 2E and 2F). Fat cells in males are observed to be different from those of females. Male fat cells contain much larger lipid droplets, reduced and less granular cytoplasm, and smaller nucleoli than female fat cells (Figure 3).

**Figure 3.**
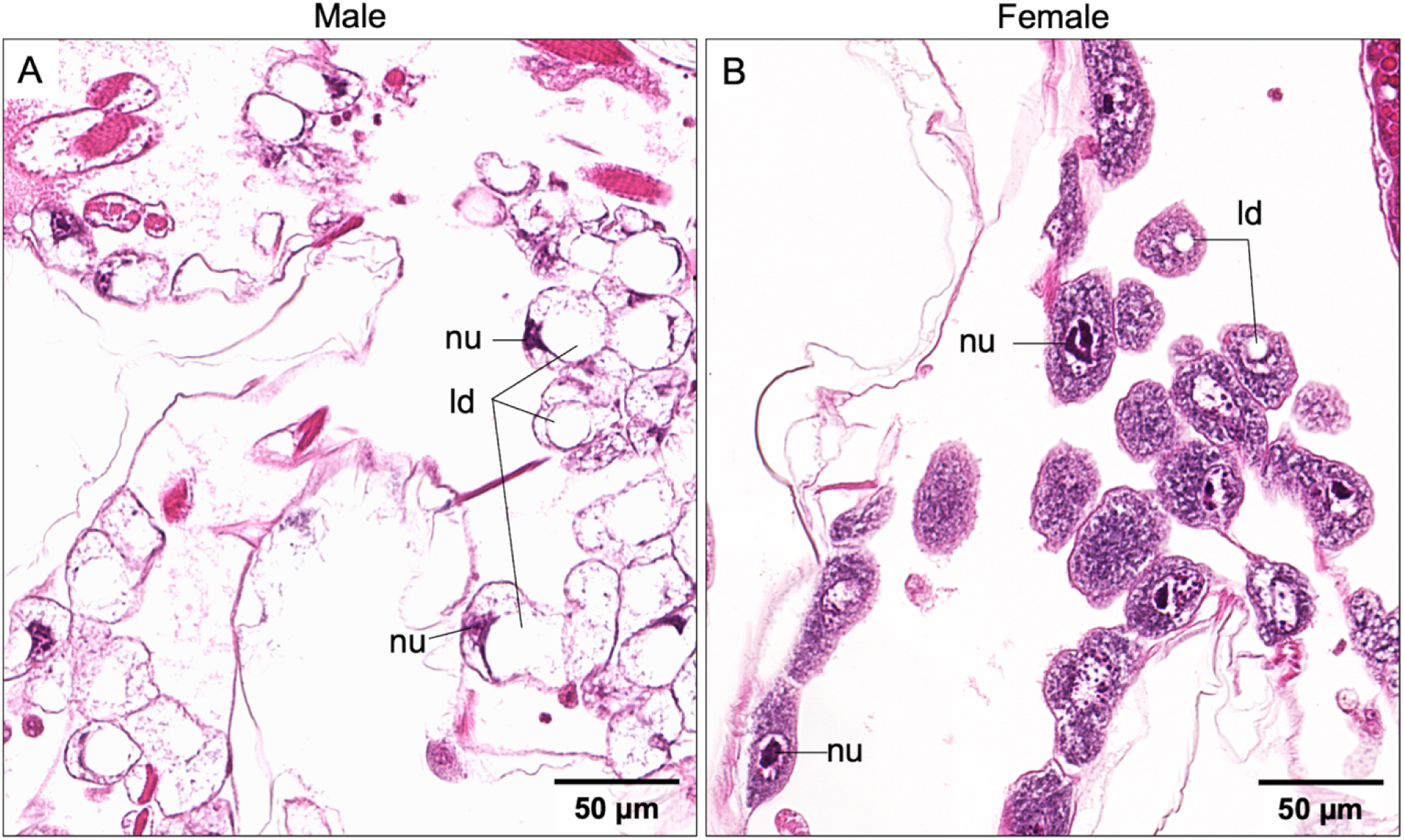
Comparison of male and female fat cells. **(A)** Fat cells in males contain larger lipid droplets (ld), reduced and less granular cytoplasm with smaller nucleoli (nu) situated at the cell periphery compared with those of females. **(B)** Fat cells in females have more granular cytoplasm with smaller lipid droplets (ld) and bigger nucleoli (nu) that often appeared subdivided.

##### Circulatory system

*Daphnia* have an open circulatory system and a myogenic heart (Stein et al., 1966; Steinsland, 1982). Since *Daphnia* are semi-transparent the beating of their hearts is easily visualized in live animals. Hemolymph (blood-like fluid) containing hemocytes (Auld et al., 2010; Metschnikoff, 1884) (Figure 2C inset) is pumped through the body cavity. In line with th literature, we also observe that the *Daphnia* heart has a pair of ostia anterior to the brood chamber, between the midgut and the dorsum (Figures 2B, C, and F). Hemoglobin is synthesized in fat cells and epithelial cells on epipodites of the thoracic limbs (Goldmann et al., 1999).

##### Digestive system

*Daphnia* are filter feeders. Food particles are filtered through filter plates consisting of setae on thoracic limbs 3 and 4, passed through maxillules and mandibles into the esophagus, which is the first part of the digestive system (Figures 2A and B). The digestive system also consists of paired hepatic ceca, midgut, and hindgut (Figures 2A-C, 2E-F) that are lined with epithelial cells and microvilli, with the columnar epithelial cells in the midgut, and the cuboidal cells in hepatic ceca and hindgut (Quaglia et al., 1976; Schultz and Kennedy, 1976). The labrum houses labral glands that have been suggested to be involved in food ingestion and endocrine function (Zaffagnini and Zeni, 1987; Zeni and Franchini, 1990) (Figures 2B-D).

##### Excretory system

The maxillary gland, also known as the shell gland, is the organ of excretion, housed between the inner and outer walls of the carapace (Smirnov, 2013). It consists of an end sac, a series of tubules, and an opening in the anterior brood chamber (Figures 2A, C, D, and F).

##### Locomotive and osmoregulation system

The swimming antennae are *Daphnia’s* primary organ of locomotion (Fryer, 1991). Each of the paired swimming antennae has a protopodite, two rami bearing setae (Agar, 1950) (Figures 2C, D, and F), and antennal muscles. *Daphnia* have five thoracic limbs (Benzie, 2005) (Figures 2B, D, and E) internal to the carapace. Movements of thoracic limbs produce a constant current that brings food particles into the digestive tract and facilitates osmotic regulation mediated by the epipodite on each thoracic limb (Kikuchi, 1983). First thoracic limbs in males are different from those of female *Daphnia*; only the male has chitinized copulatory hooks (Figure 2F) and longer setae (Figure 2E).

##### Muscular system

The muscular system occupies a significant portion of the body (Binder, 1931; Fryer, 1991). The largest muscles are the ventral and dorsal longitudinal muscles that extend along the gut, three paired antennal muscles, transverse mandibular muscles, transverse muscles of mandibles, posterior rotator of the mandibles, carapace adductor muscles, followed by groups of muscles that allow the motion of thoracic limbs and postabdomen (Figure 2). Other small muscles include those around the compound eye, labrum, and esophagus (Consi et al., 1987). All muscles are striated and surrounded by sarcoplasm, that contains many nuclei and is mostly vacuolated. Sarcoplasm is particularly abundant and more vacuolated in the antennal muscles. Male antennules also have internal muscle fibers (Figure 2D) that appear to be absent in females.

##### Nervous, sensory, and vision systems

*Daphnia* have a pigmented compound eye consisting of 22 ommatidia (Figure 2B) and a light-sensing, pigmented nauplius eye or ocellus with four lens-like bodies (Weiss et al., 2012) (Figure 2F’). Each ommatidium contains eight retinular cells sending a parallel bundle of axons, collectively as the optic nerve into the optic lobe, which is then connected to the cerebral ganglia (Figure 2B). From the cerebral ganglia, two chains of nerve cords run along the thorax, underneath the gut, and to other anatomical structures (Kress et al., 2016; McCoole et al., 2011) (Figures 2A, E, and F). Both sexes have a pair of antennules bearing a group of 9 olfactory setae or aesthetascs (Klann and Stollewerk, 2017; Rieder, 1987) but the male antennules are more prominent and elongated, uniquely fitted with a flagellum at each tip (Figure 2D’).

##### Reproductive system

The ovaries in females are paired tubular structures ending in oviducts (Figure 2A). *Daphnia* are generally cyclical parthenogens, which means that sexual (meiotic) and clonal (ameiotic) reproduction alternate (Decaestecker et al., 2009). Under favorable environmental conditions, females produce parthenogenetic eggs that are genetically identical to themselves. During clonal reproduction, oogenesis occurs in clusters of four oocytes each. Only one definitive oocyte in each set accumulates yolk granules and lipid droplets during maturation while the others become nurse cells (Rossi, 1980) (Figure 4A). After maturation, parthenogenetic eggs are released into the brood chamber through the oviducts. Fully developed, free-swimming juveniles are extruded after 3 to 4 days. Sexual reproduction is cued by environmental changes such as shorter light photoperiod, lower temperature, and over-crowding, which triggers the parthenogenetic production of genetically identical males for mating with receptive females, the endpoint being two embryos that enter a state of diapause. Unlike parthenogenetic embryos, the development of these resting embryos arrests at the 3000-cell count and enters dormancy (Chen et al., 2018). The resting embryos are encased in a chitinous shell called an ephippium that protects them from harsh environmental conditions (Figure 4C), including freezing and desiccation. Dormancy in *Daphnia* can be exceptionally long, lasting decades and even centuries (Cáceres, 1998; Mergeay et al., 2004). The resting embryos hatch when cued by favorable environmental conditions.

**Figure 4.**
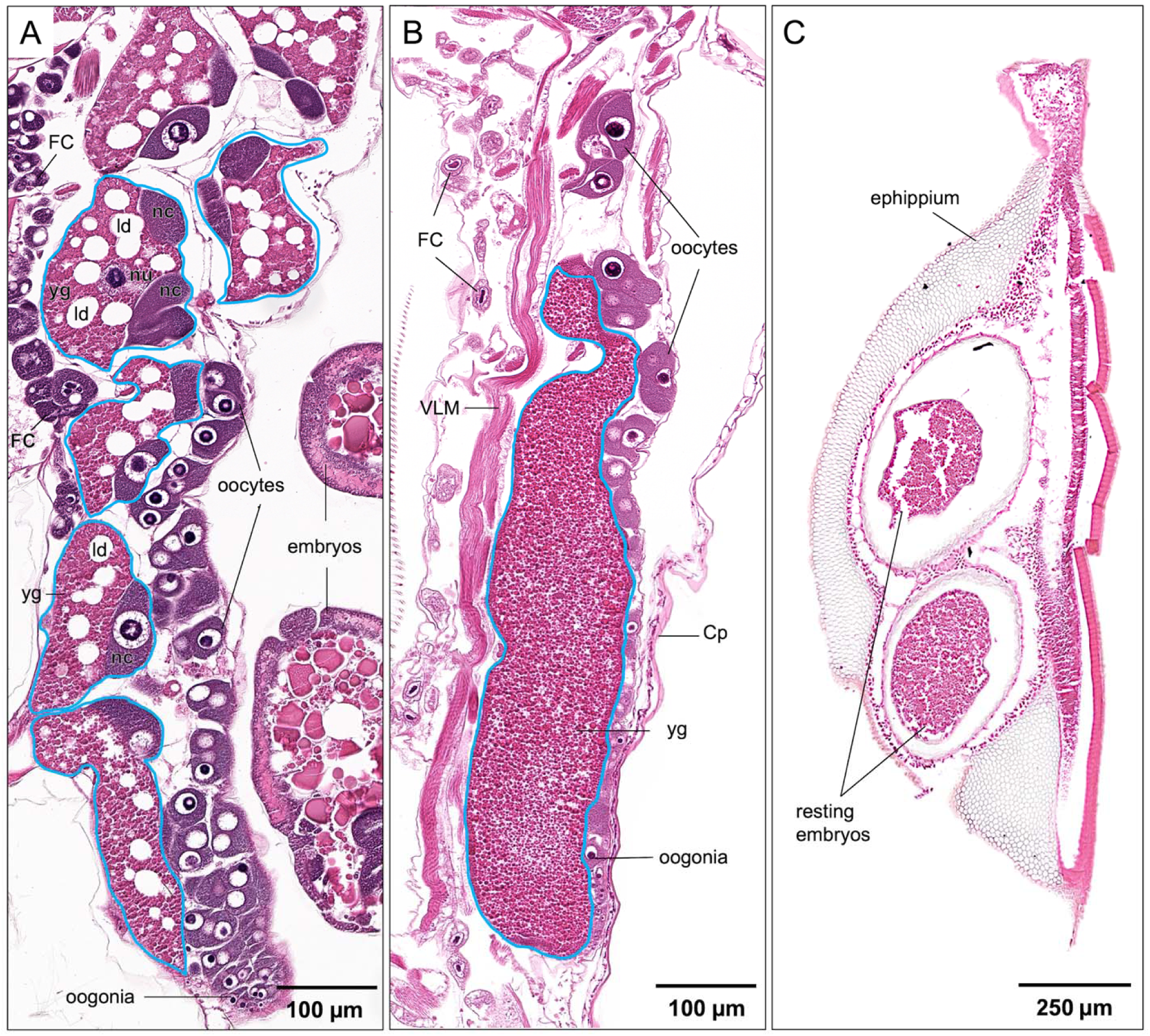
Comparison of parthenogenetic and sexual eggs. **(A)** The parthenogenetic egg contain a large amount of lipid droplets (ld) and yolk granules (yg). **(B)** The sexual eggs contain a large proportion of fine yolk granules without lipid droplets. **(C)** Resting embryos encased in the ephippium. The top embryo shows artifact. Cp, carapace; FC, fat cell; nc, nurse cell; nu, the nucleus of oocyte; VLM, ventral longitudinal muscle. A solid blue circle indicates an individual egg.

*Daphnia* testes consist of two tubular structures connected by sperm ducts to gonopores or ejaculatory openings (Figures 2D and E). Spermatogenesis begins at the testes’ walls, and mature spermatozoa are displaced inward toward the central region of the testes (Wuerz et al., 2017).

Fat cells are polyploid (Beaton and Hebert, 1989) and consist of a massive portion of lipid and glycogen (Zaffagnini and Zeni, 1986). They are typically found along the trunk, around ovaries or testes, and on the epipodites of the thoracic limbs (Figure 2). In females, these cells have a cytoplasm rich in RNA, one or several lipid droplets of various size, and one large nucleus within which a nucleolus of irregular shape resides (Zaffagnini and Zeni, 1986). The nucleolus often appears subdivided into two or more parts. They are most likely sites of vitellogenin synthesis (Zaffagnini and Zeni, 1986). Compared to female fat cells, male fat cells contain a much larger lipid droplet, reduced and less granular cytoplasm, and a smaller nucleus that is usually situated at the cell periphery (Figure 3).

#### 3.1.3 *D. magna* developmental stages

*Daphnia* embryos develop in the brood chamber before being extruded. Embryos can also develop outside of the brood chamber, in the culture medium or distilled water. *Daphnia* embryogenesis is usually staged based on the time of development after oviposition (Green, 1956; Gulbrandsen and Johnsen, 1990; Threlkeld, 1979; Toyota et al., 2016), and morphological landmarks (Mittmann et al., 2014). Due to the difficulties in orienting and sectioning the minute individual *Daphnia* embryos, DaHRA presents a selection of images of embryos found in gravid adults. The stages of the embryos were determined under the dissecting microscope according to the developmental events used by Toyota et al (2016) before the adult female were processed for histology sectioning. Developmental events corresponding with developmental stages are Stage 1: egg membrane is intact; Stage 2: egg chorion breaks down and eye spots are not yet visible; Stage 3: embryo bears two small pink or red eyes; Stage 4: embryo bears two brown or black eyes, and Stage 5: embryo bears a single median black eye. To-date, DaHRA showcases histology images of developmental stages 1, 3, 4, and 5.

At around 20°C, embryos develop within 3 days in the brood chamber after the oviposition. In our histology images, Stage 1 embryo contains yolk granules, lipid droplets, and peripheral cytoplasm (Figure 5A). During Stage 3, eye spots start to develop, and embryo will bear 2 pink eyes towards the end of Stage 3. Labrum, gut, thoracic limbs, and swimming antennae becoming distinguishable at this stage (Figure 5B). Eye pigment increases at Stage 4, and the embryo will bear two brown or black pigment cells (Figure 5C). Ocellus, cerebral ganglia and optic lobe are now distinguishable. Structures of the digestive system, such as mandibles, hepatic ceca, esophagus, midgut and hindgut can also be identified. Segments of thoracic limbs such as epipodites and exopodites are distinctly visible. Stage 5 embryo bears a single median eye, and all major anatomical structures are distinguishable as the body continues to elongate (Figure 5D). Currently, we have not been able to provide images for developmental Stage 2 which is a very short stage (about 4 hours).

**Figure 5.**
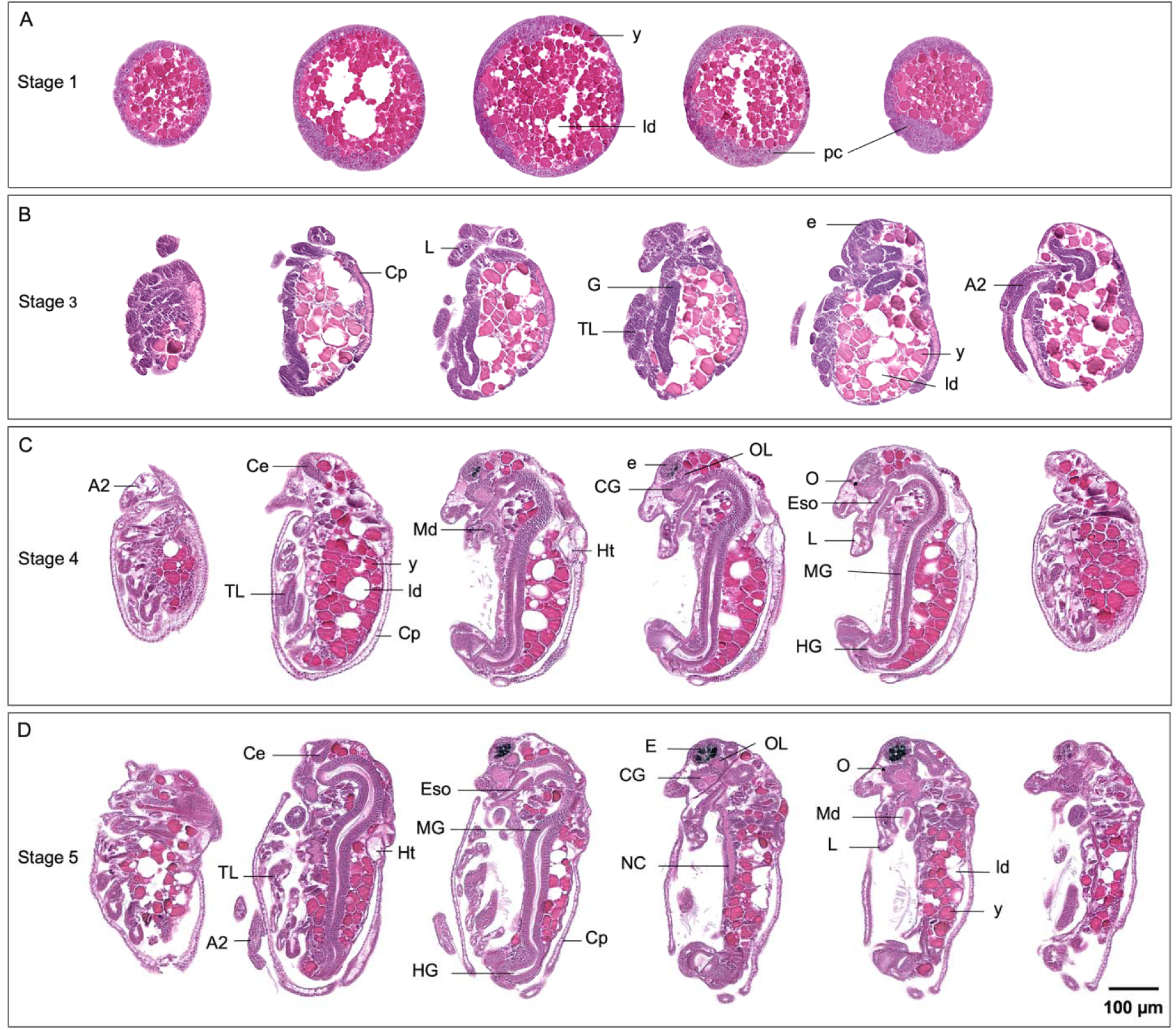
Representative developmental stages of parthenogenetic female *D. magna* in sagittal plane. **(A)** Stage 1 embryo in spherical form containing yolks (y), lipid droplets (ld), and peripheral cytoplasm (pc). Intact chorion or egg membrane is not visible here due to sectioning artifact. **(B)** Stage 3 embryo bears two pink eyes (e) with differentiation of thoracic limbs (TL), labrum (L), gut (G) and swimming antennae (A2). **(C)** Stage 4 embryo bears two brown or black eyes (e) with ocellus (O), optic lobe (OL) and cerebral ganglia (CG) being distinguishable. The labrum (L) is elongated, and segments of digestive tract are distinct. **(D)** Stage 5 embryo bears a single black median eye (E), and the body continues to elongate. A2, swimming antennae; Ce, hepatic cecum; Cp, carapace; Eso, esophagus; HG, hindgut; Md, mandible, MG, midgut; NC, nerve chord.

#### 3.1.4 *D. magna* clones

According to the diversity panel curated by Ebert research group, there are more than 200 clones of *D. magna* being cultured in the laboratories around the world (https://evolution.unibas.ch/ebert/research/referencepanel/). Clonal variation refers to the different genotypes of the same species, and they may differ strongly in their toxicological responses (Barata et al., 2002; Barber et al., 1990; Kim et al., 2023). While DaHRA showcases annotated histology from a commercial clone, we have also included histology images from a clone provided by the University of Birmingham, UK (UOB_LRV0_1) to demonstrate how this atlas can be expanded for the addition of new image datasets (http://daphnia.io/anatomy/clone/). This clone was revived from a biological archive of Lake Ring, a shallow lake in Denmark (Cuenca Cambronero et al., 2018). The clone had been sequenced to generate the first chromosomal-level genome assembly which 33,950 genes and 31,336 proteins had been annotated (Chaturvedi et al., 2023).

#### 3.1.5 An example of *D. magna* histopathology

To demonstrate how our atlas facilitates visualization of abnormal tissue architecture in the context of the whole organism, we included acetaminophen-exposed *D. magna* that show strong histopathological tissue phenotypes. “Comparison” tool shows web-based comparisons at any magnification between the exposed and to the control animals in an experiment (See Figure S5, Supplementary Material; http://daphnia.io/anatomy/treatments/).

Besides its therapeutic effects, acetaminophen is known to induce toxicological outcomes if overdose is taken over a period of time (Perananthan et al., 2024; Yoon et al., 2016). It is also a contaminant found in surface waters and wastewater throughout the world (Phong Vo et al., 2019). In our study, gravid *D. magna* were exposed to 5, 15, 25, 35, 50 µg/mL of acetaminophen for 72 hrs to allow the evaluation on both adults and developing embryos. *D. magna* exposed to 5 µg/mL acetaminophen showed no distinct changes compared to unexposed controls. Exposure to 25 µg/mL and 35 µg/mL acetaminophen caused similar phenotypes, but exposure to 50 µg/mL acetaminophen caused death by the end of the exposure. This exposure is not intended for reporting toxicological effects of acetaminophen; therefore, no replicate and error had been generated. *D. magna* exposed to 15 and 25 µg/mL acetaminophen were used to illustrate histopathological phenotypes in a whole-organism context.

Histology of the exposed *D. magna* revealed morphological alterations in various organs and tissue types. Excessive vacuolation was observed in the labral glands of *D. magna* exposed to both 15 and 25 µg/mL acetaminophen (Figures 6B and C). Besides the absence of a peritrophic membrane and partially digested food particles, the midgut and hindgut of the exposed *D. magna* showed degeneration in the epithelium lining (Figures 6E and F), with the degree being particularly conspicuous in the 25 µg/mL-exposed *D. magna* (Figure 6F). Extrusion and sloughing of epithelial cells were also observed (Figures 6E and F). Cytoplasmic swelling was noted in the midgut and hindgut epithelial cells and fat cells (Figures 6H and I).

**Figure 6.**
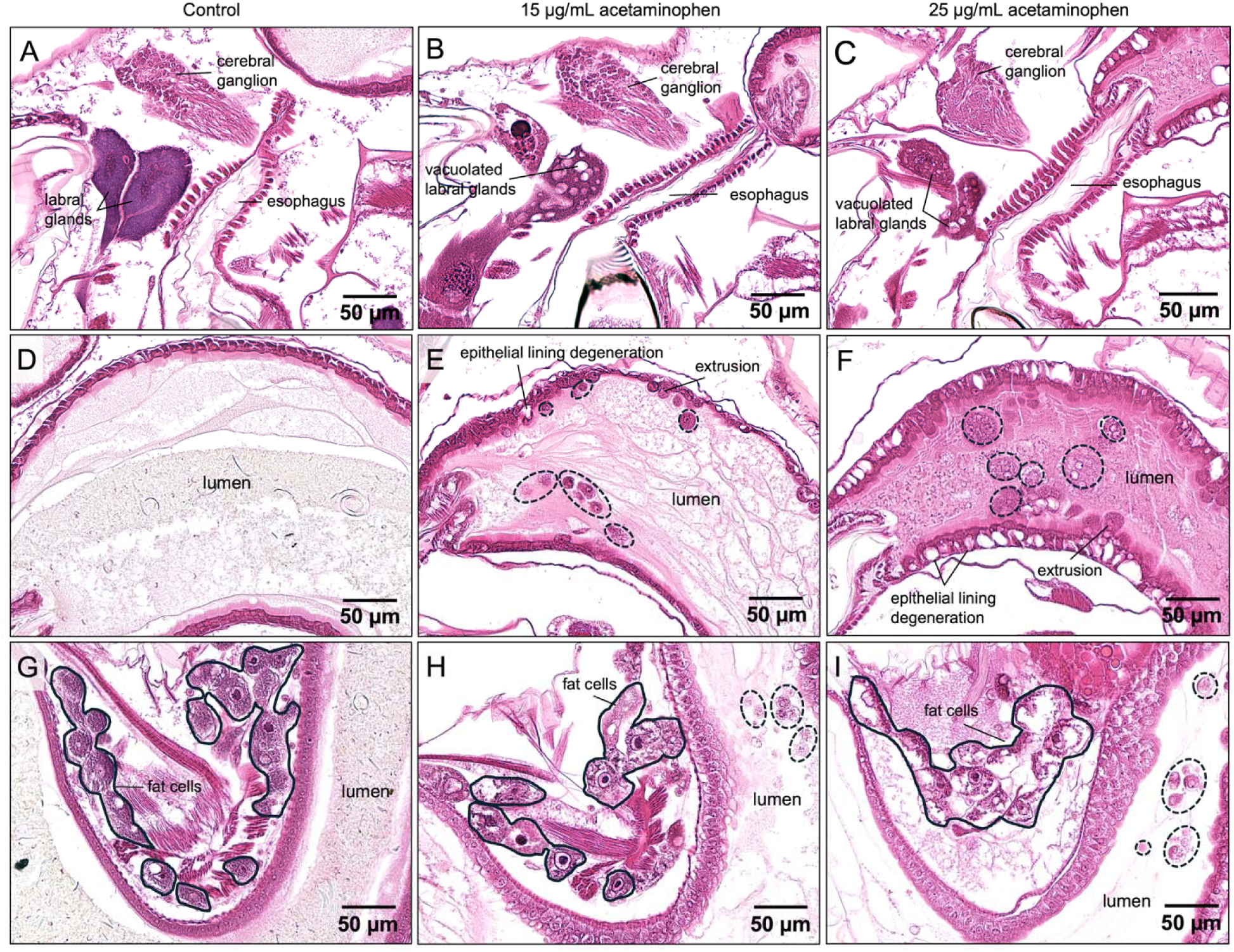
Histopathologic features of the digestive system and fat cells in *D. magna* exposed to 15 and 25 µg/mL of acetaminophen for 72 hrs. Compared to the control **(A)**, vacuolization of labral glands were present in exposed *D. magna* **(B, C)**. Extrusion, and sloughing of degenerated epithelial cells (dotted circles) was observed in exposed animals **(E, F)**, with the degree of degeneration being particularly conspicuous in the epithelium lining of 25 µg/mL- exposed *D. magna* **(F)**. Cytoplasmic swelling was observed in the fat cells and the hindgut epithelial lining of exposed *D. magna* **(H, I)**. Sloughed epithelial cells (dotted circles) were prominent in the hindgut of the exposed *D. magna* **(H, I)**.

A dead, and deformed embryos were found in the brood chamber of 25 µg/mL acetaminophen-exposed *D. magna* after 72 hrs of exposure. The deformed embryos remained in the chorion, showed development of the compound eye and gut precursors, but no visible elongation of body length, or development of the swimming antennae and thoracic limbs were evident after 72 hrs of exposure (Figure 7).

**Figure 7.**
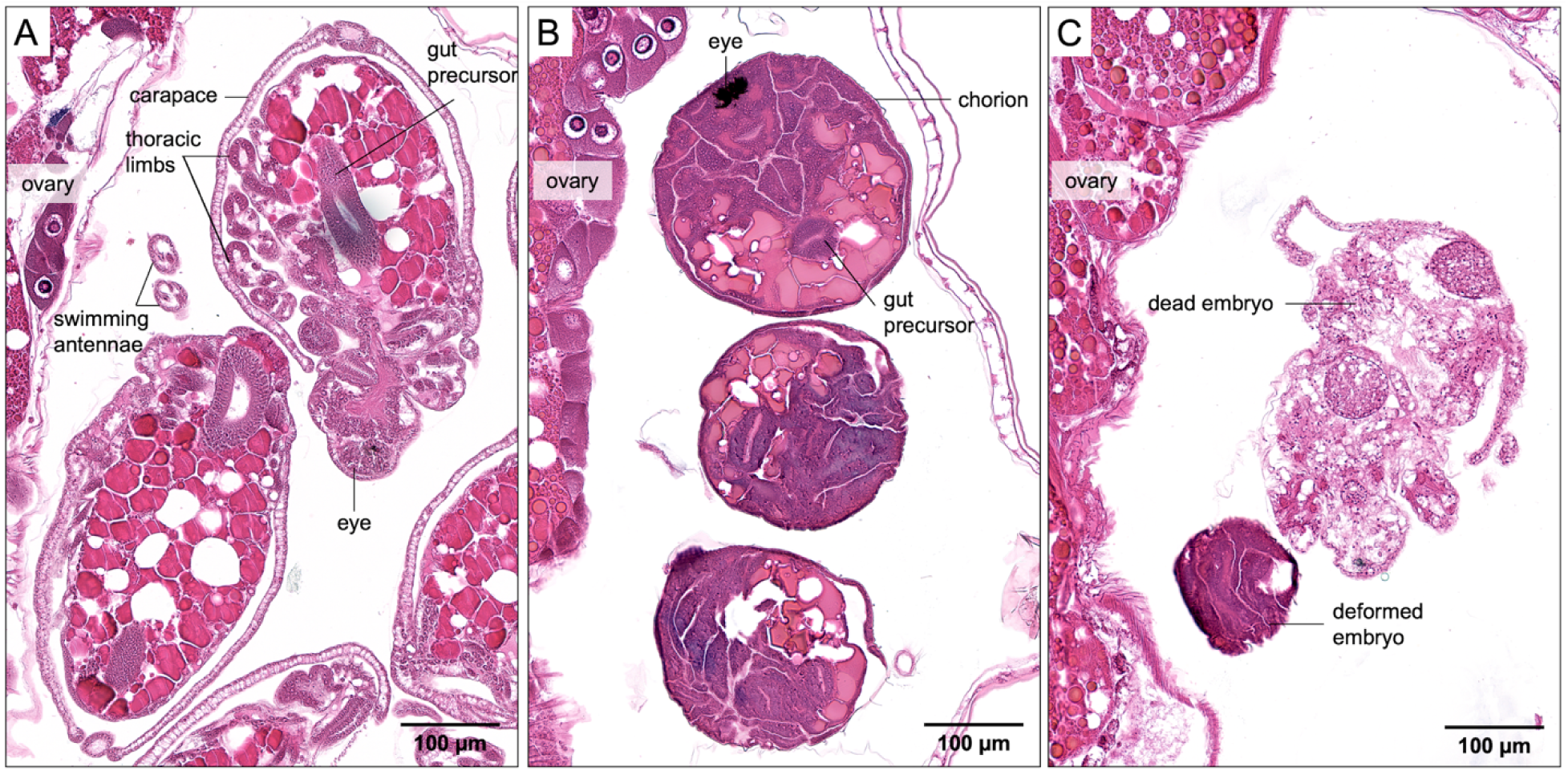
Embryotoxicity in the *D. magna* exposed to 25 µg/mL acetaminophen. **(A)** Normal embryos in the brood chamber of control compared to **(B)** deformed embryos remained in the chorion, showing some development of the compound eye and gut precursors but no visible development of the swimming antennae and thoracic limbs. **(C)** A dead embryo was also observed in the brood chamber of an exposed *D. magna*.

Toxicological effects are often quantified through apical endpoints (e.g. immobilization) in acute exposures (Organisation for Economic Co-operation and Development, 2004; United States Environmental Protection Agency, 2002) and fitness-linked life history traits in chronic exposures (Organisation for Economic Co-operation and Development, 2018; United States Environmental Protection Agency, 1996). Few studies employ ultrastructural analysis of a target organ, usually the midgut, for the pathological assessment of chemical toxicity (Bacchetta et al., 2017; Bodar et al., 1990; Heinlaan et al., 2011; Yang et al., 2010). Acetaminophen exposure performed for this work was not intended for reporting the toxicological responses. However, the observation of embryotoxicity and histopathological change in multiple tissue types (fat cells and labral glands), including the whole digestive tract (ceca, midgut, and hindgut) in the exposed *D. magna* suggested the important role of non-targeted toxicity assessment for the complete detection of phenotypes across organ systems and during embryonic development. A combination of complete histopathological phenotyping in whole *Daphnia* with toxicological omics data (e.g., transcriptomics and metabolomics) and tissue-specific biomarkers (e.g., single cell and spatial transcriptomics) will enable a comprehensive evaluation of toxicological effects and reveal tissue-specific toxicity (Tian et al., 2024). Establishing these links is essential for biomolecular data to be regulatory relevant because hazards are classified by phenotypic outcomes (behavioral, physiological, and/or histopathological). Such integration will play an important role in discovering and applying adverse outcome pathways for next-generation risk assessment (United States Environmental Protection Agency, 2014) where links between biomarkers of adversity, causative agents, and organ-specific effects remain to be well-established. Using the principle of evolutionary and functional conservation of genes and pathways in organisms across the tree of life, hypotheses on targets of toxicity can be extrapolated across non-target species, including humans (“The Precision Toxicology initiative,” 2023).

### 3.2 Future directions

We have created an annotated, interactive atlas for web-based access to *D. magna* microanatomy using a single commercially available clone of *D. magna* to present reasonable examples of what “normal” microanatomy looks like. Clonal and intraspecific variation are important components in *Daphnia* biology (Ho et al., 2019; Miyakawa et al., 2015; Tolardo et al., 2016) where differential responses to toxicants and other stressors have been reported for physiological and reproduction parameters (Barata et al., 2002; Barber et al., 1990; Kim et al., 2023), but remain to be explored with regard to histopathological change and tissue-specific phenotypes.

A key aspect of this work is our commitment to open science, enabling broader participation of *Daphnia* community to utilize and further develop this atlas. A clone UOB_LRV0_1 has been included in the current phase of DaHRA, and through collaboration and additional resources, we anticipate this atlas to include additional datasets possibly covering clonal and/or intraspecific genetic variation, examples of how intraspecific variation influences tissue-specific phenotypes, and most importantly, observation of other histopathological change. It is our hope for this atlas to be an informational tool to the *Daphnia* community, but it is not intended and should not be utilized as “control” for chemical-exposure and toxicity-testing experiments.

Web-based atlases have served as a platform for the systematic integration of spatial and molecular data (Asp et al., 2019; Snyder et al., 2019; Thul and Lindskog, 2018; Yao et al., 2023). As integrative tissue-based or spatial atlases for human and mammalian models are constantly being improved , DaHRA is envisioned as a potential integrative platform for anchoring *Daphnia* multi-omics data with tissue phenotypes. This will increase its potential value as a tool for exploring the spatial and chemical complexities of biological systems.

In summary, our first, open-source visualization platform for *D. magna* microanatomy provides a baseline knowledge of *Daphnia*’s cells, extracellular tissue, and their arrangements in health as a foundation for the detection of aberrant tissue phenotypes. Its user-friendly interface and global web-based access will facilitate broader contributions to (eco)toxicology, biology and beyond.

## Supplementary Material (in order of citation)

Table S1. Fixatives and fixation parameters tested for best preservation of whole *D. magna* samples

Figure S1. “Ballooning” artifact observed in *D. magna* sample fixed with PFA and NBF

Figure S2. Comparison of histological sections generated with Bouin’s, PFA and NBF

Figure S3. Agarose embedding of *D. mag*na samples using agarose block with triangular wells

Figure S4. Anatomy of adult female and male *D. magna*

File S1. *Daphnia* anatomy glossary

Table S2. Tissue processing steps for serial dehydration and infiltration of *D. magna* samples with Formula R paraffin in tissue processor

Table S3. Automated steps for staining *D. magna* 5-μm sections with Harris’ hematoxylin and eosin in an auto-stainer

Table S4. Anatomical ontology for DaHRA

Figure S5. Overview of DaHRA displaying annotated histopathological data

## Acknowledgments

The authors thank Debra Shearer and Chadwick Harris for sectioning the paraffin blocks and staining the slides. The authors thank Dr. Matthew Lanza, Department of Comparative Medicine, Penn State College of Medicine for confirming the histopathology in the exposed *D. magna* samples. The authors also thank Patrick Leibich and the Department of Surgery, Penn State College of Medicine for 3D-SLA printing the casting molds. The authors appreciate Jessica Christ’s help with the graphical abstract. This work was supported by the Penn State Human Health and Environment Seed Grant funded by Pennsylvania Department of Health Commonwealth Universal Research Enhancement Program Grant (to KCA), the National Institutes of Health grant 1R24OD18559 to KCC, and the Jake Gittlen Laboratories for Cancer Research gift fund. The Department of Health specifically disclaims responsibility for any analyses, interpretations, or conclusions. This work contributes to the PrecisionTox project (https://precisiontox.org/) that received funding from the European Union’s Horizon 2020 research and innovation program under grant agreement No 965406. This output reflects only the authors’ views, and the European Union cannot be held responsible for any use that may be made of the information contained therein.

## Author Contributions

Mee Ngu: Data curation, Investigation, Methodology, Project administration, Visualization, Writing-original draft; Daniel Vanselow: Data curation, Visualization, Software; Carolyn Zaino: Data curation, Visualization; Alex Lin: Methodology; Jean Copper: Methodology; Margaret Beaton: Validation; Luisa Orsini: Conceptualization; John Colbourne: Conceptualization; Keith Cheng: Conceptualization, Funding Acquisition, Resources; Khai Ang: Conceptualization, Funding Acquisition, Investigation, Methodology, Resources, Supervision. All authors contributed to Writing-Review and Editing.

## References

Abdullahi, M., Li, X., Abdallah, M.A.-E., Stubbings, W., Yan, N., Barnard, M., Guo, L.-H., Colbourne, J.K., Orsini, L., 2022. Daphnia as a sentinel species for environmental health protection: a perspective on biomonitoring and bioremediation of chemical pollution. Environ. Sci. Technol. 56, 14237–14248. 10.1021/acs.est.2c01799

Agar, W.E., 1950. The swimming setae of Daphnia carinata. J. Cell Sci. s3-91, 353–368. 10.1242/jcs.s3-91.16.353

Armit, C., Richardson, L., Venkataraman, S., Graham, L., Burton, N., Hill, B., Yang, Y., Baldock, R.A., 2017. eMouseAtlas: An atlas-based resource for understanding mammalian embryogenesis. Dev. Biol. 423, 1–11. 10.1016/j.ydbio.2017.01.023

Asp, M., Giacomello, S., Larsson, L., Wu, C., Fürth, D., Qian, X., Wärdell, E., Custodio, J., Reimegård, J., Salmén, F., Österholm, C., Ståhl, P.L., Sundström, E., Åkesson, E., Bergmann, O., Bienko, M., Månsson-Broberg, A., Nilsson, M., Sylvén, C., Lundeberg, J., 2019. A Spatiotemporal Organ-Wide Gene Expression and Cell Atlas of the Developing Human Heart. Cell 179, 1647–1660.e19. 10.1016/j.cell.2019.11.025

Auld, S.K.J.R., Scholefield, J.A., Little, T.J., 2010. Genetic variation in the cellular response of Daphnia magna (Crustacea: Cladocera) to its bacterial parasite. Proc. R. Soc. B Biol. Sci. 277, 3291–3297. 10.1098/rspb.2010.0772

Bacchetta, R., Santo, N., Marelli, M., Nosengo, G., Tremolada, P., 2017. Chronic toxicity effects of ZnSO4 and ZnO nanoparticles in Daphnia magna. Environ. Res. 152, 128–140. 10.1016/j.envres.2016.10.006

Barata, C., Baird, D.J., Soares, A.M.V.M., 2002. Determining genetic variability in the distribution of sensitivities to toxic stress among and within field populations of Daphnia magna. Environ. Sci. Technol. 36, 3045–3049. 10.1021/es0158556

Barber, I., Baird, D.J., Calow, P., 1990. Clonal variation in general responses of Daphnia magna Straus to toxic stress. II. Physiological effects. Funct. Ecol. 4, 409–414. 10.2307/2389603

Beaton, M.J., Hebert, P.D.N., 1989. Miniature genomes and endopolyploidy in cladoceran crustaceans. Genome 32, 1048–1053. 10.1139/g89-552

Bednarska, A., 2006. Adaptive changes in morphology of Daphnia filter appendages in response to food stress. Polish Journal of Ecology 54, 663–668.

Benzie, J.A.H., 2005. The Genus Daphnia (including Daphniopsis) (Anomopoda:Daphniidae). Kenobi Productions.

Binder, G., 1931. Das Muskelsystem von Daphnia. Int. Rev. Gesamten Hydrobiol. Hydrogr. 26, 54–98. 10.1002/iroh.19310260104

Bodar, C.W.M., van Donselaar, E.G., Herwig, H.J., 1990. Cytopathological investigations of digestive tract and storage cells in Daphnia magna exposed to cadmium and tributyltin. Aquat. Toxicol. 17, 325–337. 10.1016/0166-445X(90)90015-H

Byeon, E., Kim, M.-S., Kim, D.-H., Lee, Y., Jeong, H., Lee, Jin-Sol, Hong, S.-A., Park, J.C., Kang, H.-M., Sayed, A.E.-D.H., Kato, Y., Bae, S., Watanabe, H., Lee, Y.H., Lee, Jae-Seong, 2022. The freshwater water flea Daphnia magna NIES strain genome as a resource for CRISPR/Cas9 gene targeting: The glutathione S-transferase omega 2 gene. Aquat. Toxicol. 242, 106021. 10.1016/j.aquatox.2021.106021

Cáceres, C.E., 1998. Interspecific variation in the abundance, production, and emergence of Daphnia diapausing eggs. Ecology 79, 1699–1710. 10.2307/176789

Campos, B., Fletcher, D., Piña, B., Tauler, R., Barata, C., 2018. Differential gene transcription across the life cycle in Daphnia magna using a new all genome custom-made microarray. BMC Genomics 19, 370. 10.1186/s12864-018-4725-7

Cardinale, B.J., Duffy, J.E., Gonzalez, A., Hooper, D.U., Perrings, C., Venail, P., Narwani, A., Mace, G.M., Tilman, D., Wardle, D.A., Kinzig, A.P., Daily, G.C., Loreau, M., Grace, J.B., Larigauderie, A., Srivastava, D.S., Naeem, S., 2012. Biodiversity loss and its impact on humanity. Nature 486, 59–67. 10.1038/nature11148

Chaturvedi, A., Li, X., Dhandapani, V., Marshall, H., Kissane, S., Cuenca-Cambronero, M., Asole, G., Calvet, F., Ruiz-Romero, M., Marangio, P., Guigó, R., Rago, D., Mirbahai, L., Eastwood, N., Colbourne, J.K., Zhou, J., Mallon, E., Orsini, L., 2023. The hologenome of Daphnia magna reveals possible DNA methylation and microbiome-mediated evolution of the host genome. Nucleic Acids Res. 51, 9785–9803. 10.1093/nar/gkad685

Chen, L., Barnett, R.E., Horstmann, M., Bamberger, V., Heberle, L., Krebs, N., Colbourne, J.K., Gómez, R., Weiss, L.C., 2018. Mitotic activity patterns and cytoskeletal changes throughout the progression of diapause developmental program in Daphnia. BMC Cell Biol. 19, 30. 10.1186/s12860-018-0181-0

Christensen, A.K., Owusu, N.G., Jean-Louis, D., 2018. Carapace epithelia are rich in large filamentous actin bundles in Daphnia magna, Daphnia pulex, and Sida crystallina (Crustacea: Cladocera). Invertebr. Biol. 137, 49–59. 10.1111/ivb.12204

Consi, T.R., Macagno, E.R., Necles, N., 1987. The oculomotor system of Daphnia magna. The eye muscles and their motor neurons. Cell Tissue Res. 247, 515–523. 10.1007/BF00215744

Copper, J.E., Budgeon, L.R., Foutz, C.A., van Rossum, D.B., Vanselow, D.J., Hubley, M.J., Clark, D.P., Mandrell, D.T., Cheng, K.C., 2018. Comparative analysis of fixation and embedding techniques for optimized histological preparation of zebrafish. Comp. Biochem. Physiol. Toxicol. Pharmacol. CBP 208, 38–46. 10.1016/j.cbpc.2017.11.003

Cuenca Cambronero, M., Marshall, H., De Meester, L., Davidson, T.A., Beckerman, A.P., Orsini, L., 2018. Predictability of the impact of multiple stressors on the keystone species Daphnia. Sci. Rep. 8, 17572. 10.1038/s41598-018-35861-y

Cuenca-Cambronero, M., Pantel, J.H., Marshall, H., Nguyen, T.T.T., Tomero-Sanz, H., Orsini, L., 2021. Evolutionary mechanisms underpinning fitness response to multiple stressors in Daphnia. Evol. Appl. 14, 2457–2469. 10.1111/eva.13258

de Oliveira, L.L.D., Antunes, S.C., Gonçalves, F., Rocha, O., Nunes, B., 2016. Acute and chronic ecotoxicological effects of four pharmaceuticals drugs on cladoceran Daphnia magna. Drug Chem. Toxicol. 39, 13–21. 10.3109/01480545.2015.1029048

Decaestecker, E., De Meester, L., Mergeay, J., 2009. Cyclical parthenogenesis in Daphnia: sexual versus asexual reproduction, in: Schön, I., Martens, K., Dijk, P. (Eds.), Lost Sex: The Evolutionary Biology of Parthenogenesis. Springer Netherlands, Dordrecht, pp. 295–316. 10.1007/978-90-481-2770-2_15

Du, J., Mei, C.-F., Ying, G.-G., Xu, M.-Y., 2016. Toxicity thresholds for diclofenac, acetaminophen and ibuprofen in the water flea Daphnia magna. Bull. Environ. Contam. Toxicol. 97, 84–90. 10.1007/s00128-016-1806-7

Ebert, D., 2022. Daphnia as a versatile model system in ecology and evolution. EvoDevo 13, 16. 10.1186/s13227-022-00199-0

Ebert, D., 2005. Introduction to Daphnia Biology, Ecology, Epidemiology, and Evolution of Parasitism in Daphnia. National Center for Biotechnology Information (US).

Edwards, C., 1980. The anatomy of Daphnia mandibles. Trans. Am. Microsc. Soc. 99, 2–24. 10.2307/3226076

European Chemicals Agency, 2020. The use of alternatives to testing on animals for the REACH Regulation.

Fraga, N., Benito, D., Briaudeau, T., Izagirre, U., Ruiz, P., 2022. Toxicopathic effects of lithium in mussels. Chemosphere 307, 136022. 10.1016/j.chemosphere.2022.136022

Fryer, G., 1991. Functional morphology and the adaptive radiation of the Daphniidae (Branchiopoda: Anomopoda). Philos. Trans. R. Soc. Lond. B. Biol. Sci. 331, 1–99. 10.1098/rstb.1991.0001

Fuller, R., Landrigan, P.J., Balakrishnan, K., Bathan, G., Bose-O’Reilly, S., Brauer, M., Caravanos, J., Chiles, T., Cohen, A., Corra, L., Cropper, M., Ferraro, G., Hanna, J., Hanrahan, D., Hu, H., Hunter, D., Janata, G., Kupka, R., Lanphear, B., Lichtveld, M., Martin, K., Mustapha, A., Sanchez-Triana, E., Sandilya, K., Schaefli, L., Shaw, J., Seddon, J., Suk, W., Téllez-Rojo, M.M., Yan, C., 2022. Pollution and health: a progress update. Lancet Planet. Health 6, e535–e547. 10.1016/S2542-5196(22)00090-0

Goldmann, T., Becher, B., Wiedorn, K.H., Pirow, R., Deutschbein, M.E., Vollmer, E., Paul, R.J., 1999. Epipodite and fat cells as sites of hemoglobin synthesis in the branchiopod crustacean Daphnia magna. Histochem. Cell Biol. 112, 335–339. 10.1007/pl00007905

Graham, E., Moss, J., Burton, N., Armit, C., Richardson, L., Baldock, R., 2015. The atlas of mouse development eHistology resource. Development 142, 1909–1911. 10.1242/dev.124917

Green, J., 1956. Growth, Size and Reproduction in Daphnia (crustacea: Cladocera). Proc. Zool. Soc. Lond. 126, 173–204. 10.1111/j.1096-3642.1956.tb00432.x

Gulbrandsen, J., Johnsen, G.H., 1990. Temperature-dependent development of parthenogenetic embryos in Daphnia pulex de Geer. J. Plankton Res. 12, 443–453. 10.1093/plankt/12.3.443

Halcrow, K., 1976. The fine structure of the carapace integument of Daphnia magna Straus (Crustacea Branchiopoda). Cell Tissue Res. 169, 267–276. 10.1007/BF00214213

Harrill, J.A., Viant, M.R., Yauk, C.L., Sachana, M., Gant, T.W., Auerbach, S.S., Beger, R.D., Bouhifd, M., O’Brien, J., Burgoon, L., Caiment, F., Carpi, D., Chen, T., Chorley, B.N., Colbourne, J., Corvi, R., Debrauwer, L., O’Donovan, C., Ebbels, T.M.D., Ekman, D.R., Faulhammer, F., Gribaldo, L., Hilton, G.M., Jones, S.P., Kende, A., Lawson, T.N., Leite, S.B., Leonards, P.E.G., Luijten, M., Martin, A., Moussa, L., Rudaz, S., Schmitz, O., Sobanski, T., Strauss, V., Vaccari, M., Vijay, V., Weber, R.J.M., Williams, A.J., Williams, A., Thomas, R.S., Whelan, M., 2021. Progress towards an OECD reporting framework for transcriptomics and metabolomics in regulatory toxicology. Regul. Toxicol. Pharmacol. RTP 125, 105020. 10.1016/j.yrtph.2021.105020

Hebert, P.D.N., Ward, R.D., 1972. Inheritance during parthenogenesis in Daphnia magna. Genetics 71, 639–642.

Heinlaan, M., Kahru, A., Kasemets, K., Arbeille, B., Prensier, G., Dubourguier, H.-C., 2011. Changes in the Daphnia magna midgut upon ingestion of copper oxide nanoparticles: A transmission electron microscopy study. Water Res. 45, 179–190. 10.1016/j.watres.2010.08.026

Hines, A., Staff, F.J., Widdows, J., Compton, R.M., Falciani, F., Viant, M.R., 2010. Discovery of metabolic signatures for predicting whole organism toxicology. Toxicol. Sci. Off. J. Soc. Toxicol. 115, 369–378. 10.1093/toxsci/kfq004

Hiruta, C., Tochinai, S., 2014. Formation and structure of the ephippium (resting egg case) in relation to molting and egg laying in the water flea Daphnia pulex De Geer (Cladocera: Daphniidae). J. Morphol. 275, 760–767. 10.1002/jmor.20255

Ho, E.K.H., Macrae, F., Latta, L.C., IV, Benner, M.J., Sun, C., Ebert, D., Schaack, S., 2019. Intraspecific Variation in Microsatellite Mutation Profiles in Daphnia magna. Mol. Biol. Evol. 36, 1942–1954. 10.1093/molbev/msz118

Huang, J., Wang, Q., Liu, S., Zhang, M., Liu, Y., Sun, L., Wu, Y., Tu, W., 2021. Crosstalk between histological alterations, oxidative stress and immune aberrations of the emerging PFOS alternative OBS in developing zebrafish. Sci. Total Environ. 774, 145443. 10.1016/j.scitotenv.2021.145443

Jankowski, M.D., Fairbairn, D.J., Baller, J.A., Westerhoff, B.M., Schoenfuss, H.L., 2022. Using the Daphnia magna transcriptome to distinguish water source: wetland and stormwater case studies. Environ. Toxicol. Chem. 41, 2107–2123. 10.1002/etc.5392

Joshy, A., Sharma, S.R.K., Mini, K.G., Gangadharan, S., Pranav, P., 2022. Histopathological evaluation of bivalves from the southwest coast of India as an indicator of environmental quality. Aquat. Toxicol. 243, 106076. 10.1016/j.aquatox.2022.106076

Kikuchi, S., 1983. The fine structure of the gill epithelium of a fresh-water flea, Daphnia magna (Crustacea: Phyllopoda) and changes associated with acclimation to various salinities. I. Normal fine structure. Cell Tissue Res. 229, 253–268. 10.1007/BF00214974

Kim, H.J., Koedrith, P., Seo, Y.R., 2015. Ecotoxicogenomic approaches for understanding molecular mechanisms of environmental chemical toxicity using aquatic invertebrate, Daphnia model organism. Int. J. Mol. Sci. 16, 12261–12287. 10.3390/ijms160612261

Kim, J., Coutellec, M.-A., Lee, S., Choi, J., 2023. Insights into the mechanisms of within-species variation in sensitivity to chemicals: A case study using daphnids exposed to CMIT/MIT biocide. Ecotoxicol. Environ. Saf. 258, 114967. 10.1016/j.ecoenv.2023.114967

Klann, M., Stollewerk, A., 2017. Evolutionary variation in neural gene expression in the developing sense organs of the crustacean Daphnia magna. Dev. Biol. 424, 50–61. 10.1016/j.ydbio.2017.02.011

Kress, T., Harzsch, S., Dircksen, H., 2016. Neuroanatomy of the optic ganglia and central brain of the water flea Daphnia magna (Crustacea, Cladocera). Cell Tissue Res. 363, 649–677. 10.1007/s00441-015-2279-4

Landrigan, P.J., Fuller, R., Acosta, N.J.R., Adeyi, O., Arnold, R., Basu, N. (Nil), Baldé, A.B., Bertollini, R., Bose-O’Reilly, S., Boufford, J.I., Breysse, P.N., Chiles, T., Mahidol, C., Coll-Seck, A.M., Cropper, M.L., Fobil, J., Fuster, V., Greenstone, M., Haines, A., Hanrahan, D., Hunter, D., Khare, M., Krupnick, A., Lanphear, B., Lohani, B., Martin, K., Mathiasen, K.V., McTeer, M.A., Murray, C.J.L., Ndahimananjara, J.D., Perera, F., Potočnik, J., Preker, A.S., Ramesh, J., Rockström, J., Salinas, C., Samson, L.D., Sandilya, K., Sly, P.D., Smith, K.R., Steiner, A., Stewart, R.B., Suk, W.A., Schayck, O.C.P. van, Yadama, G.N., Yumkella, K., Zhong, M., 2018. The Lancet Commission on pollution and health. The Lancet 391, 462–512. 10.1016/S0140-6736(17)32345-0

Lee, B.-Y., Choi, B.-S., Kim, M.-S., Park, J.C., Jeong, C.-B., Han, J., Lee, J.-S., 2019. The genome of the freshwater water flea Daphnia magna: A potential use for freshwater molecular ecotoxicology. Aquat. Toxicol. 210, 69–84. 10.1016/j.aquatox.2019.02.009

Majno, G., Joris, I., 2004. Cells, tissues, and disease : principles of general pathology, 2nd ed. ed. New York : Oxford University Press.

Manjunatha, B., Seo, E., Bangyappagari, D., Lee, S.J., 2022. Histopathological and ultrastructural alterations reveal the toxicity of particulate matter (PM2.5) in adult zebrafish. J. Hazard. Mater. Adv. 7, 100135. 10.1016/j.hazadv.2022.100135

McCoole, M.D., Baer, K.N., Christie, A.E., 2011. Histaminergic signaling in the central nervous system of Daphnia and a role for it in the control of phototactic behavior. J. Exp. Biol. 214, 1773–1782. 10.1242/jeb.054486

Mergeay, J., Verschuren, D., Kerckhoven, L.V., Meester, L.D., 2004. Two hundred years of a diverse Daphnia community in Lake Naivasha (Kenya): effects of natural and human-induced environmental changes. Freshw. Biol. 49, 998–1013. 10.1111/j.1365-2427.2004.01244.x

Metschnikoff, E., 1884. A disease of Daphnia caused by a yeast. A contribution to the theory of phagocytes as agents for attack on disease-causing organisms. Archiv. Pathol. Anat. Physiol. Klin. Med 96, 177–195.

Miner, B.E., De Meester, L., Pfrender, M.E., Lampert, W., Hairston, N.G., 2012. Linking genes to communities and ecosystems: Daphnia as an ecogenomic model. Proc. R. Soc. B Biol. Sci. 279, 1873–1882. 10.1098/rspb.2011.2404

Mittmann, B., Ungerer, P., Klann, M., Stollewerk, A., Wolff, C., 2014. Development and staging of the water flea Daphnia magna (Straus, 1820; Cladocera, Daphniidae) based on morphological landmarks. EvoDevo 5, 12. 10.1186/2041-9139-5-12

Miyakawa, H., Sugimoto, N., Kohyama, T.I., Iguchi, T., Miura, T., 2015. Intra-specific variations in reaction norms of predator-induced polyphenism in the water flea Daphnia pulex. Ecol. Res. 30, 705–713. 10.1007/s11284-015-1272-4

Naidu, R., Biswas, B., Willett, I.R., Cribb, J., Kumar Singh, B., Paul Nathanail, C., Coulon, F., Semple, K.T., Jones, K.C., Barclay, A., Aitken, R.J., 2021. Chemical pollution: A growing peril and potential catastrophic risk to humanity. Environ. Int. 156, 106616. 10.1016/j.envint.2021.106616

Organisation for Economic Co-operation and Development, 2018. Daphnia magna reproduction test (OECD TG 211). OECD. 10.1787/9789264304741-12-en

Organisation for Economic Co-operation and Development, 2004. Test No. 202: Daphnia sp. acute immobilisation test. OECD.

Orsini, L., Gilbert, D., Podicheti, R., Jansen, M., Brown, J.B., Solari, O.S., Spanier, K.I., Colbourne, J.K., Rusch, D.B., Decaestecker, E., Asselman, J., De Schamphelaere, K.A.C., Ebert, D., Haag, C.R., Kvist, J., Laforsch, C., Petrusek, A., Beckerman, A.P., Little, T.J., Chaturvedi, A., Pfrender, M.E., De Meester, L., Frilander, M.J., 2016. Daphnia magna transcriptome by RNA-Seq across 12 environmental stressors. Sci. Data 3, 160030. 10.1038/sdata.2016.30

Palmer, J.A., Smith, A.M., Gryshkova, V., Donley, E.L.R., Valentin, J.-P., Burrier, R.E., 2020. A targeted metabolomics-based assay using human induced pluripotent stem cell-derived cardiomyocytes identifies structural and functional cardiotoxicity potential. Toxicol. Sci. 174, 218–240. 10.1093/toxsci/kfaa015

Perananthan, V., Shihana, F., Chiew, A.L., George, J., Dawson, A., Buckley, N.A., 2024. Intestinal injury in paracetamol overdose (ATOM-8). J. Gastroenterol. Hepatol. 39, 920– 926. 10.1111/jgh.16450

Phong Vo, H.N., Le, G.K., Hong Nguyen, T.M., Bui, X.-T., Nguyen, K.H., Rene, E.R., Vo, T.D.H., Thanh Cao, N.-D., Mohan, R., 2019. Acetaminophen micropollutant: Historical and current occurrences, toxicity, removal strategies and transformation pathways in different environments. Chemosphere 236, 124391. 10.1016/j.chemosphere.2019.124391

Quaglia, A., Sabelli, B., Villani, L., 1976. Studies on the intestine of Daphnidae (Crustacea, Cladocera) ultrastructure of the midgut of Daphnia magna and Daphnia obtusa. J. Morphol. 150, 711–725. 10.1002/jmor.1051500306

Ramírez-Duarte, W., Rondon Barragan, I., Eslava-Mocha, P., 2008. Acute toxicity and histopathological alterations of Roundup® herbicide on “cachama blanca” (Piaractus brachypomus). Pesqui. Veterinária Bras. 28, 547. 10.1590/S0100-736X2008001100002

Rieder, N., 1987. The ultrastructure of the so-called olfactory setae on the antennula of Daphnia magna Straus (Crustacea, Cladocera), in: Forró, L., Frey, D.G. (Eds.), Cladocera, Developments in Hydrobiology. Springer Netherlands, Dordrecht, pp. 175–181. 10.1007/978-94-009-4039-0_20

Rossi, F., 1980. Comparative observations on the female reproductive system and parthenogenetic oogenesis in Cladocera. Bolletino Zool. 47, 21–38. 10.1080/11250008009440317

Santana, L.M.B.M., Damasceno, É.P., Loureiro, S., Soares, A.M.V.M., Pousão-Ferreira, P., Abessa, D.M.S., Martins, R., Pavlaki, M.D., 2023. An easy-to-use histological technique for small biological samples of Senegalese Sole larvae. Appl. Sci. 13, 2346. 10.3390/app13042346

Schindelin, J., Arganda-Carreras, I., Frise, E., Kaynig, V., Longair, M., Pietzsch, T., Preibisch, S., Rueden, C., Saalfeld, S., Schmid, B., Tinevez, J.-Y., White, D.J., Hartenstein, V., Eliceiri, K., Tomancak, P., Cardona, A., 2012. Fiji: an open-source platform for biological-image analysis. Nat. Methods 9, 676–682. 10.1038/nmeth.2019

Schneider, C.A., Rasband, W.S., Eliceiri, K.W., 2012. NIH Image to ImageJ: 25 years of image analysis. Nat. Methods 9, 671–675. 10.1038/nmeth.2089

Schultz, T.W., Kennedy, J.R., 1976. The fine structure of the digestive system of Daphnia pulex (Crustacea: Cladocera). Tissue Cell 8, 479–490. 10.1016/0040-8166(76)90008-2

Shaw, J.R., Pfrender, M.E., Eads, B.D., Klaper, R., Callaghan, A., Sibly, R.M., Colson, I., Jansen, B., Gilbert, D., Colbourne, J.K., 2008. Daphnia as an emerging model for toxicological genomics, in: Hogstrand, C., Kille, P. (Eds.), Advances in Experimental Biology, Comparative Toxicogenomics. Elsevier, pp. 165–328. 10.1016/S1872-2423(08)00005-7

Smirnov, N.N., 2013. Physiology of the Cladocera. Physiol. Cladocera 1–336.

Snyder, M.P., Lin, S., Posgai, A., Atkinson, M., Regev, A., Rood, J., Rozenblatt-Rosen, O., Gaffney, L., Hupalowska, A., Satija, R., Gehlenborg, N., Shendure, J., Laskin, J., Harbury, P., Nystrom, N.A., Silverstein, J.C., Bar-Joseph, Z., Zhang, K., Börner, K., Lin, Y., Conroy, R., Procaccini, D., Roy, A.L., Pillai, A., Brown, M., Galis, Z.S., Cai, L., Shendure, J., Trapnell, C., Lin, S., Jackson, D., Snyder, M.P., Nolan, G., Greenleaf, W.J., Lin, Y., Plevritis, S., Ahadi, S., Nevins, S.A., Lee, H., Schuerch, C.M., Black, S., Venkataraaman, V.G., Esplin, E., Horning, A., Bahmani, A., Zhang, K., Sun, X., Jain, S., Hagood, J., Pryhuber, G., Kharchenko, P., Atkinson, M., Bodenmiller, B., Brusko, T., Clare-Salzler, M., Nick, H., Otto, K., Posgai, A., Wasserfall, C., Jorgensen, M., Brusko, M., Maffioletti, S., Caprioli, R.M., Spraggins, J.M., Gutierrez, D., Patterson, N.H., Neumann, E.K., Harris, R., deCaestecker, M., Fogo, A.B., van de Plas, R., Lau, K., Cai, L., Yuan, G.-C., Zhu, Q., Dries, R., Yin, P., Saka, S.K., Kishi, J.Y., Wang, Y., Goldaracena, I., Laskin, J., Ye, D., Burnum-Johnson, K.E., Piehowski, P.D., Ansong, C., Zhu, Y., Harbury, P., Desai, T., Mulye, J., Chou, P., Nagendran, M., Bar-Joseph, Z., Teichmann, S.A., Paten, B., Murphy, R.F., Ma, J., Kiselev, V.Yu., Kingsford, C., Ricarte, A., Keays, M., Akoju, S.A., Ruffalo, M., Gehlenborg, N., Kharchenko, P., Vella, M., McCallum, C., Börner, K., Cross, L.E., Friedman, S.H., Heiland, R., Herr, B., Macklin, P., Quardokus, E.M., Record, L., Sluka, J.P., Weber, G.M., Nystrom, N.A., Silverstein, J.C., Blood, P.D., Ropelewski, A.J., Shirey, W.E., Scibek, R.M., Mabee, P., Lenhardt, W.C., Robasky, K., Michailidis, S., Satija, R., Marioni, J., Regev, A., Butler, A., Stuart, T., Fisher, E., Ghazanfar, S., Rood, J., Gaffney, L., Eraslan, G., Biancalani, T., Vaishnav, E.D., Conroy, R., Procaccini, D., Roy, A., Pillai, A., Brown, M., Galis, Z., Srinivas, P., Pawlyk, A., Sechi, S., Wilder, E., Anderson, J., HuBMAP Consortium, Writing Group, Caltech-UW TMC, Stanford-WashU TMC, UCSD TMC, University of Florida TMC, Vanderbilt University TMC, California Institute of Technology TTD, Harvard TTD, Purdue TTD, Stanford TTD, HuBMAP Integration, V., and Engagement (HIVE) Collaboratory: Carnegie Mellon, Tools Component, Harvard Medical School, T.C., Indiana University Bloomington, M.C., Pittsburgh Supercomputing Center and University of Pittsburgh, I. and E.C., University of South Dakota, C.C., New York Genome Center, M.C., NIH HuBMAP Working Group, 2019. The human body at cellular resolution: the NIH Human Biomolecular Atlas Program. Nature 574, 187–192. 10.1038/s41586-019-1629-x

Stein, R.J., Richter, W.R., Zussman, R.A., Brynjolfsson, G., 1966. Ultrastructural characterization of Daphnia heart muscle. J. Cell Biol. 29, 168–170.

Steinsland, A.J., 1982. Heart ultrastructure in Daphnia pulex De Geer (Crustacea, Branchiopoda, Cladocera). J. Crustac. Biol. 2, 54–58. 10.2307/1548112

Stoks, R., Govaert, L., Pauwels, K., Jansen, B., De Meester, L., 2016. Resurrecting complexity: the interplay of plasticity and rapid evolution in the multiple trait response to strong changes in predation pressure in the water flea Daphnia magna. Ecol. Lett. 19, 180–190. 10.1111/ele.12551

Stollewerk, A., 2010. The water flea Daphnia - a “new” model system for ecology and evolution? J. Biol. 9, 21. 10.1186/jbiol212

Stucki, A.O., Barton-Maclaren, T.S., Bhuller, Y., Henriquez, J.E., Henry, T.R., Hirn, C., Miller-Holt, J., Nagy, E.G., Perron, M.M., Ratzlaff, D.E., Stedeford, T.J., Clippinger, A.J., 2022. Use of new approach methodologies (NAMs) to meet regulatory requirements for the assessment of industrial chemicals and pesticides for effects on human health. Front. Toxicol. 4. 10.3389/ftox.2022.964553

The Precision Toxicology initiative, 2023. . Toxicol. Lett. 383, 33–42. 10.1016/j.toxlet.2023.05.004

Threlkeld, S.T., 1979. Estimating cladoceran birth rates: The importance of egg mortality and the egg age distribution. Limnol. Oceanogr. 24, 601–612. 10.4319/lo.1979.24.4.0601

Thul, P.J., Lindskog, C., 2018. The human protein atlas: A spatial map of the human proteome. Protein Sci. Publ. Protein Soc. 27, 233–244. 10.1002/pro.3307

Tian, R., Guan, M., Chen, L., Wan, Y., He, L., Zhao, Z., Gao, T., Zong, L., Chang, J., Zhang, J., 2024. Mechanism insights into the histopathological changes of polypropylene microplastics induced gut and liver in zebrafish. Ecotoxicol. Environ. Saf. 280, 116537. 10.1016/j.ecoenv.2024.116537

Tolardo, M., da Silva Ferrão-Filho, A., Santangelo, J.M., 2016. Species and clone-dependent effects of tilapia fish (Cichlidae) on the morphology and life-history of temperate and tropical Daphnia. Ecol. Res. 31, 333–342. 10.1007/s11284-016-1337-z

Toyota, K., Hiruta, C., Ogino, Y., Miyagawa, S., Okamura, T., Onishi, Y., Tatarazako, N., Iguchi, T., 2016. Comparative Developmental Staging of Female and Male Water Fleas Daphnia pulex and Daphnia magna During Embryogenesis. Zoolog. Sci. 33, 31–37. 10.2108/zs150116

United States Environmental Protection Agency, 2014. Next Generation Risk Assessment: incorporation of recent advances in molecular, computational, and systems biology (final report) [WWW Document]. URL https://cfpub.epa.gov/ncea/risk/recordisplay.cfm?deid=286690 (accessed 2.22.22).

United States Environmental Protection Agency, 2002. Methods for measuring the acute toxicity of effluents and receiving waters to freshwater and marine organisms.

United States Environmental Protection Agency, 1996. Ecological effects test guidelines OCSPP850.1300: Daphnid chronic toxicity test.

van der Ven, L.T.M., Wester, P.W., Vos, J.G., 2003. Histopathology as a tool for the evaluation of endocrine disruption in zebrafish (Danio rerio). Environ. Toxicol. Chem. 22, 908–913.

Walsh, M.R., Packer, M., Beston, S., Funkhouser, C., Gillis, M., Holmes, J., Goos, J., 2018. Daphnia as a Model for Eco-evolutionary Dynamics, in: Thiel, M., Wellborn, G.A. (Eds.), Life Histories: Volume 5. Oxford University Press, p. 0. 10.1093/oso/9780190620271.003.0016

Weiss, L.C., Tollrian, R., Herbert, Z., Laforsch, C., 2012. Morphology of the Daphnia nervous system: a comparative study on Daphnia pulex, Daphnia lumholtzi, and Daphnia longicephala. J. Morphol. 273, 1392–1405. 10.1002/jmor.20068

Wester, P.W., Canton, J.H., 1991. The usefulness of histopathology in aquatic toxicity studies. Comp. Biochem. Physiol. Part C Comp. Pharmacol. 100, 115–117. 10.1016/0742-8413(91)90135-G

Wuerz, M., Huebner, E., Huebner, J., 2017. The morphology of the male reproductive system, spermatogenesis and the spermatozoon of Daphnia magna (Crustacea: Branchiopoda). J. Morphol. 278, 1536–1550. 10.1002/jmor.20729

Yang, X.Y., Edelmann, R.E., Oris, J.T., 2010. Suspended C60 nanoparticles protect against short-term UV and fluoranthene photo-induced toxicity, but cause long-term cellular damage in Daphnia magna. Aquat. Toxicol., Aquatic Toxicology of Nanomaterials 100, 202–210. 10.1016/j.aquatox.2009.08.011

Yao, Z., van Velthoven, C.T.J., Kunst, M., Zhang, M., McMillen, D., Lee, C., Jung, W., Goldy, J., Abdelhak, A., Aitken, M., Baker, K., Baker, P., Barkan, E., Bertagnolli, D., Bhandiwad, A., Bielstein, C., Bishwakarma, P., Campos, J., Carey, D., Casper, T., Chakka, A.B., Chakrabarty, R., Chavan, S., Chen, M., Clark, M., Close, J., Crichton, K., Daniel, S., DiValentin, P., Dolbeare, T., Ellingwood, L., Fiabane, E., Fliss, T., Gee, J., Gerstenberger, J., Glandon, A., Gloe, J., Gould, J., Gray, J., Guilford, N., Guzman, J., Hirschstein, D., Ho, W., Hooper, M., Huang, M., Hupp, M., Jin, K., Kroll, M., Lathia, K., Leon, A., Li, S., Long, B., Madigan, Z., Malloy, J., Malone, J., Maltzer, Z., Martin, N., McCue, R., McGinty, R., Mei, N., Melchor, J., Meyerdierks, E., Mollenkopf, T., Moonsman, S., Nguyen, T.N., Otto, S., Pham, T., Rimorin, C., Ruiz, A., Sanchez, R., Sawyer, L., Shapovalova, N., Shepard, N., Slaughterbeck, C., Sulc, J., Tieu, M., Torkelson, A., Tung, H., Valera Cuevas, N., Vance, S., Wadhwani, K., Ward, K., Levi, B., Farrell, C., Young, R., Staats, B., Wang, M.-Q.M., Thompson, C.L., Mufti, S., Pagan, C.M., Kruse, L., Dee, N., Sunkin, S.M., Esposito, L., Hawrylycz, M.J., Waters, J., Ng, L., Smith, K., Tasic, B., Zhuang, X., Zeng, H., 2023. A high-resolution transcriptomic and spatial atlas of cell types in the whole mouse brain. Nature 624, 317–332. 10.1038/s41586-023-06812-z

Yoon, E., Babar, A., Choudhary, M., Kutner, M., Pyrsopoulos, N., 2016. Acetaminophen-induced hepatotoxicity: a comprehensive update. J. Clin. Transl. Hepatol. 4, 131–142. 10.14218/JCTH.2015.00052

Zaffagnini, F., Zeni, C., 1987. Ultrastructural investigations on the labral glands of Daphnia obtusa (Crustacea, Cladocera). J. Morphol. 193, 23–33. 10.1002/jmor.1051930104

Zaffagnini, F., Zeni, C., 1986. Considerations on some cytological and ultrastructural observations on fat cells of Daphnia (Crustacea, Cladocera). Bollettino di zoologia 53, 33–39. 10.1080/11250008609355480

Zeni, C., Franchini, A., 1990. A preliminary histochemical study on the labral glands of Daphnia obtusa (Crustacea, Cladocera). Acta Histochem 88, 175–181.

Zhang, L., Baer, K.N., 2000. The influence of feeding, photoperiod and selected solvents on the reproductive strategies of the water flea, Daphnia magna. Environ. Pollut. 110, 425–430. 10.1016/S0269-7491(99)00324-3

