## Supplementary Material for "A web-based histology atlas for the freshwater sentinel species *Daphnia magna*"

**Table S1**: Fixatives and fixation parameters tested for best preservation of whole *D. magna* samples

| Fixative | Fixation time and temperature | Decalcification by cold 6% formic acid time and temperature |
| --- | --- | --- |
| Bouin’s solution | 24 h, 21°C  48 h, 4°C | 24 h, 21°C  24 h, 4°C |
|  | 48 h, 21°C | 24 h, 21°C |
| 4% Paraformaldehyde | 24 h, 21°C  48 h, 4°C | 24 h, 21°C  24 h, 4°C |
|  | 48 h, 21°C | 24 h, 4°C |
| 10% Buffered Formalin Phosphate | 24 h, 4°C  48 h, 4°C | 24 h, 21°C  24 h, 4°C |
|  | 48 h, 21°C | 24 h, 4°C |

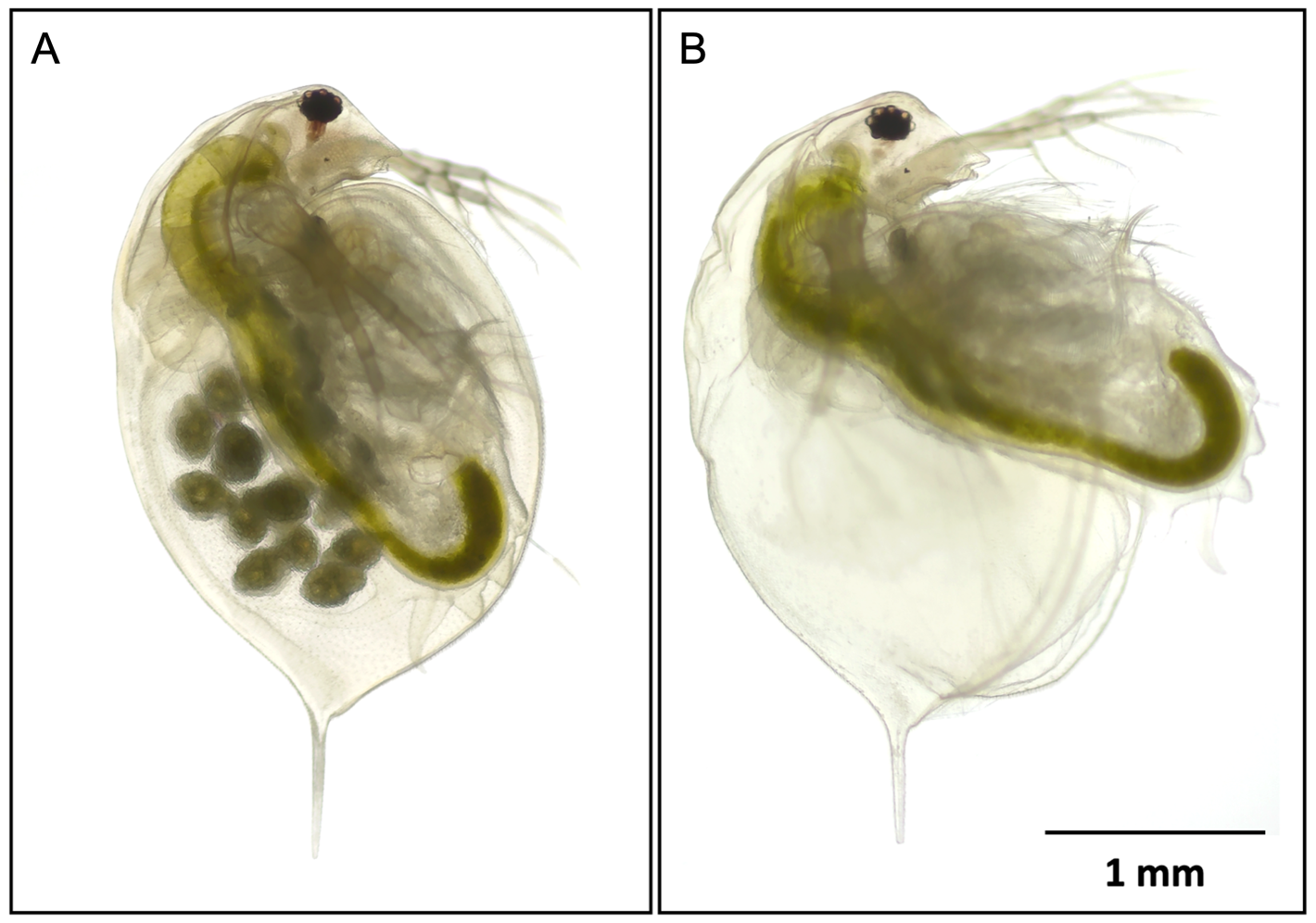

**Figure S1.** **“Ballooning” artifact observed in *D. magna* sample fixed with 4% PFA and 10% NBF.** **(A)** A well-preserved *D. magna* sample. **(B)** A *D. magna* sample with the “ballooning” artifact. This artifact is characterized by bulging of the carapace and ventral extension of the postabdomen. When both occurred, eggs or embryos were released from the brood chambers. Ballooned samples were difficult to orient during agarose pre-embedding. The mechanism of ballooning is unclear. Haney and Hall (1973) (92) suggested that it is likely related to a death-induced osmotic imbalance of the connective tissue that attached the carapace to the animal’s body. Fixation at 4 °C prevented this artifact, but it recurred after de-calcification in cold 6% formic acid at 4° C.

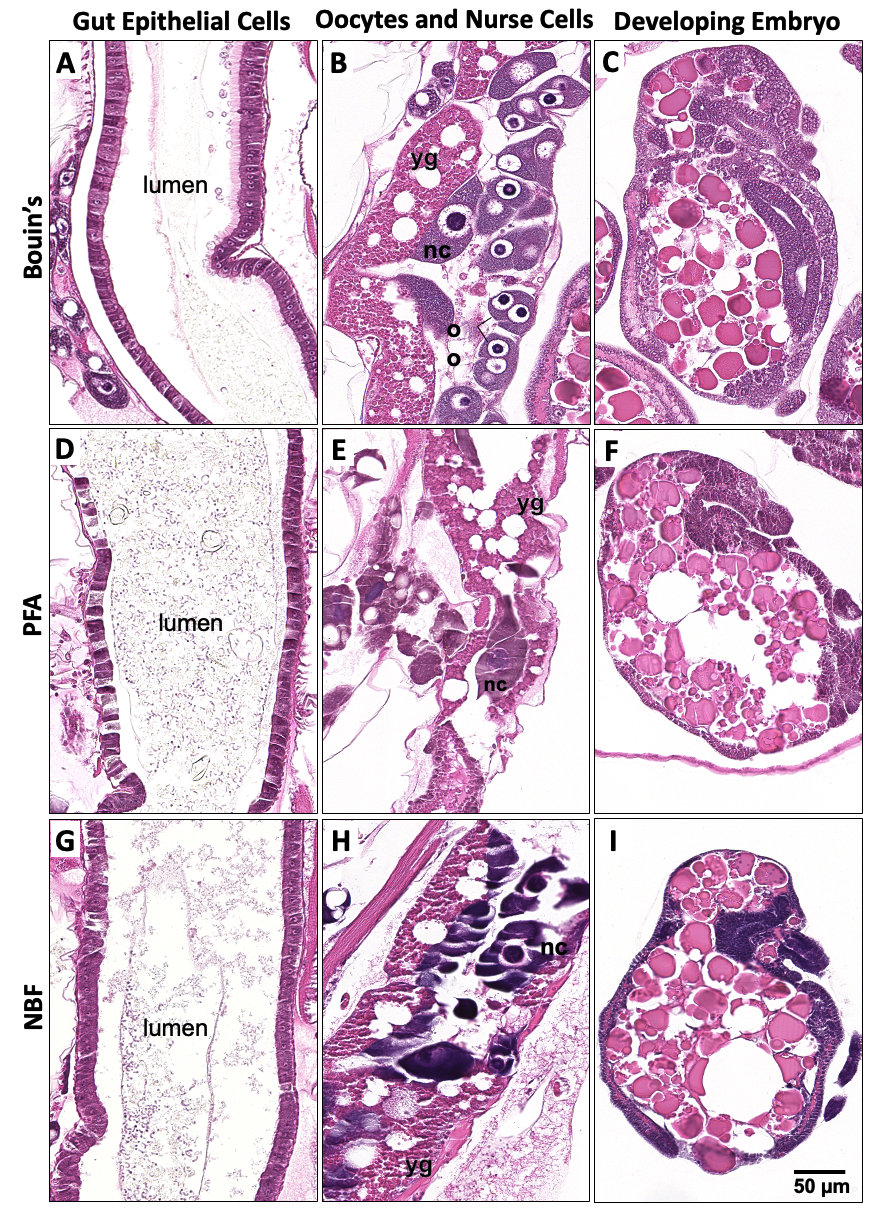

**Figure S2. Comparison of histological sections generated with different fixatives: Bouin’s, PFA and NBF.** Bouin’s fixed samples showing intact gut sections, with microvilli and epithelial cell nuclei **(A)**, as well as nurse cells (nc), oocytes (oo) and yolk granules (yg) clearly visible in the ovary **(B)**. In comparison, PFA or NBF fixed samples show less cellular clarity due to the low preservation of histological sections. Cellular details across the developing embryos are clearly visible in samples fixed with Bouin’s **(C)** as compared to other two fixatives **(F and I)**.

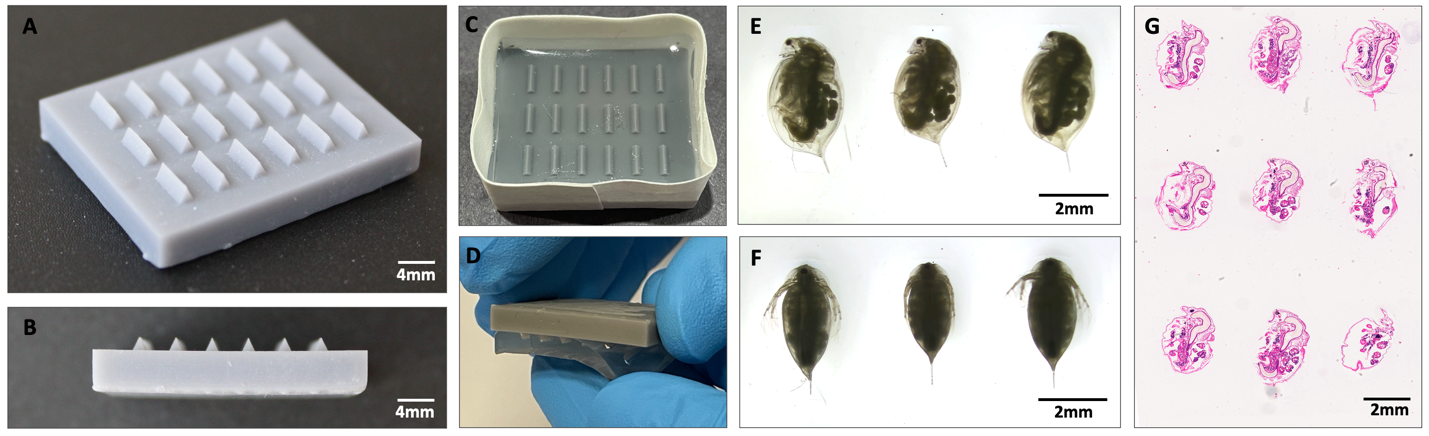

**Figure S3. Agarose embedding of *D. mag*na samples using agarose block with triangular wells.**  **(A)** Top view and **(B)** side view of mold printed by stereo-lithography (3D-SLA) at 25 µm resolution for a smooth surface allowing easy removal of agarose blocks. **(C)** Casting 1% agarose block in the taped mold. **(D)** Agarose block is removed from the mold after solidification by peeling the gel downwards. *D. magna* samples laid on their sides with a swimming antenna in the wells **(E)**, the rostra facing the same direction for sagittal plane sectioning and positioned in the wells on their back **(F)** for coronal and transverse plane sectioning. **(G)** Histological section showing the position of samples at sagittal plane. STL file for the casting mold can be downloaded from http://daphnia.io/resources/.

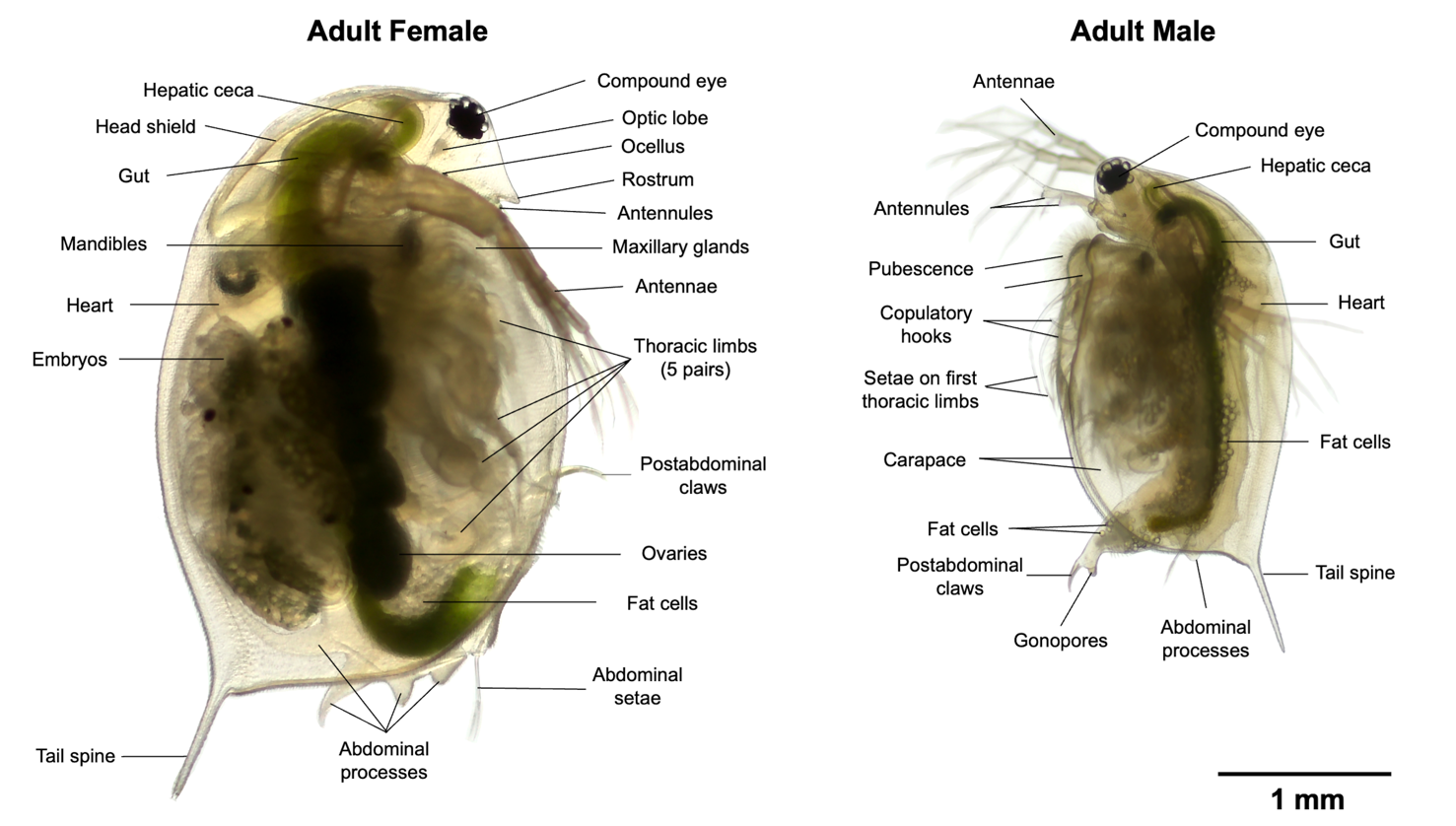

**Figure S4.** Anatomy of adult female and male *D. magna*

**File S1:** *Daphnia* anatomy glossary

**Abdominal process** (*syn. abdominal outgrowth*)

One of typically two dorsal processes projecting from border of trunk and postabdomen; considered to serve in closing off brood chamber posteriorly

### Abdominal seta *(pl. setae)*

One of two elaborate setae located on small protuberance on dorsal surface of postabdomen

### Adductor muscle (of carapace)

Muscle attached to carapace for pulling it to the body (thorax)

### Aesthetasc

One in a tuft of sensory projections at tip of each antennule

**Antenna** *(syn. second antenna, pl. antennae)*

One of the second and much larger pair of antennae; located laterally near posterior margin of head; biramous, consisting of relatively large basal part (protopodite) bearing two- to four-segmented dorsal and ventral branches (rami), serves as principle locomotory organ and moved by relatively large antennal muscles

**Antennule** *(syn. first antenna)*

One of the first and typically much smaller pair of antennae; located ventrally near posterior margin of head; uniramous, unsegmented, with tuft of terminal aesthetascs

**Anus**

Posterior opening of digestive tract at end of postabdomen

**Adductor muscle**

Muscles attached to the carapace that pull it to the body, or connect the carapace

**Brood Chamber** *(syn. brood pouch)*

Any space or sac-like cavity utilized as a uterus, in which eggs or embryos are developed

**Carapace** (*syn. shell*)

A cuticular, usually calcified, single-piece (univalved) shield which, if fully developed, covers only posterior part of body (trunk); laterally compressed, with ventral gape, and therefore occasionally described as bivalved

**Cerebral ganglion** *(syn. supraesophageal ganglion)*

One of a pair of ganglia (or fused median ganglion) situated in the head or anterior part of the body in front of or dorsal to the esophagus; often termed as brain

### Chitin

A resistant complex chemical compound, the chief constituent of the exoskeleton, or carapace

### Compound eye

One of two large photosensitive organs on head; sessile, typically fused, and enclosed within optic vesicle; movable by series of optic muscles

### Hepatic cecum (*syn. cecum, digestive cecum, midgut diverticulum*)

One of two lateral digestive organs of midgut; located in head posterior to border of esophagus and midgut

### Dorsal

Of, or pertaining to, the back or upper surface

### End sac (of maxillary gland)

Expanded proximal section of each maxillary gland; opens to exterior via tubule (convoluted duct)

**Endite**

Inwardly (medially) directed, setose lobe of basal part (protopodite) of thoracic limb; first endite typically well delimited, the following endites increasingly incorporated into endopodite

### Ephippium

Egg case formed by walls of brood pouch, typically separated from rest of carapace during molting

**Epipodite**

In thoracic limb, laterally (outwardly) directed lobe projecting from base of protopodite, represents only lobe articulated with limb and serves in respiration.

**Esophagus** *(syn. foregut, stomodeum)*

Relatively short and narrow anterior section of digestive tract, ectodermal in origin and line with cuticle; cast off with molt

**Exopodite**

Lobe-like distal branch of thoracic limb; lacks articulation with protopodal part of limb

**Exoskeleton**

Chitinous or calcified outer integument covering trunk and limbs

### Filter plate *(syn. filter comb)*

### One of the paired structures on thoracic limbs (usually third and fourth) that serves to filter suspended feeding matter

### Food groove

Elongate median groove between bases of thoracic limbs, passed anteriorly along food groove, and transferred to mouth

**Fornix**

Ridge in lateral part of cephalon above insertion of antennal muscles

### Gnathobase

Paired endites used to manipulate or move food

**Gonopore**

Opening of male reproductive system to exterior

### Head *(syn. cephalon)*

Anterior of two divisions of body (head, trunk); bearing antennules, antennae, mandibles, maxillules, (reduced) maxillae, compound and naupliar eyes, and may be covered by a head shield; not enclosed by carapace

### Head shield *(syn. cephalic shield)*

Variously developed, unpaired, shield-like structure covering head

### Heart

In circulatory system, typically short, muscular pumping organ located anterior to brood chamber in trunk; blood enters heart through single pair of ostia and is pumped anteriorly into a sinus

### Hemolymph

The circulating fluid; composed of cells and plasma; often loosely termed as blood.

**Hemocyte**

A mesodermal cell, sessile, circulating in the hemolymph; often loosely termed as blood cell

### Hindgut *(syn. proctodeum)*

Short posterior most section of digestive tract between midgut and anus

### Integument

Outer covering of exoskeleton

**Labrum** *(syn. upper lip)*

Unpaired, median lip-like structure posterior to mouth, distal ends of mandibles extend under labrum

### Levator muscle

A muscle serving to raise an organ or part

### Mandible

Paired, jaw-like cutting, grinding, or crushing appendages, consists of a fairly hard, chitinised region

### Maxillule (*syn. first maxilla, maxillula)*

Cephalic appendages immediately posterior to mandibles, serving as mouth part.

**Maxillary gland** (*syn. shell gland*)

One of two excretory organs located in the anterior region of the thorax and opening ventrally at level of maxillule; forms several loops (tubule).

**Midgut**

Elongate section of digestive tract between esophagus and hindgut

**Nerve cord**

Pair of widely separated, longitudinal nerve cords extending into trunk from posterior part of cerebral ganglion; forms ladder-like chain with relatively few, occasionally fused ganglia

**Ocellus** *(syn. median eye, naupliar eye)*

Small median photosensitive organ located ventrally on head between mouth and compound eye

**Ommatidium** *(pl. ommatidia)*

One of the component units of a compound eye, consisting essentially of an optical and a sensory part

### Optic lobe

Lateral extensions of the protocerebrum or nervous system for innervation of an eye

### Optic nerve

Nerve(s) leading from cerebral ganglion to each (fused) compound eye

**Ostium** *(pl. ostia)*

One of two lateral openings in heart; hemolymph enters heart through ostia and is pumped anteriorly

**Ovary**

Paired section of female reproductive system in which eggs are produced; typically extends through anterior part of trunk, one along each side of midgut, opens dorsally via oviduct into brood chamber

**Oviduct**

Short and narrow section of female reproductive system between posterior part of each ovary and dorsal brood chamber

**Peritropic membrane**

Chitinous membrane, secreted in anterior region of midgut and surrounding feces; considered to protect midgut lining from damage during passage of undigestible material

**Postabdomen**

Recurved (turned ventrally and forward) posteriormost region of trunk, bears pair of abdominal setae proximally and caudal rami distally), as well as series of spines (anal spines) and denticles, terminating in 2 claws

**Postabdominal claw** (*syn. claw, caudal ramus, terminal claw*)

One of two relatively short, claw-shaped projections at end of postabomen, may bear up to three series (pectens) of minute spines

**Protocerebrum**

One of two anterior dorsal cerebral ganglia; receives the optic nerves

**Protopodite** *(syn. protopod)*

Proximal part of thoracic limb; bears one or two distinguishable endites along inner margin and epipodite on outer margin, as well as distal endopodite and exopodite

Basal part of each antenna; relatively long, bearing two- to four-segmented dorsal and ventral branches (rami)

**Ramus** *(pl. rami)*

Branch of appendage, refers either to dorsal or ventral branches of antennae

**Ridge**

Elevated ridge along carapace.

**Rostrum**

Anterior (ventrally directed) beaklike extension of head

### Seta *(pl. setae)*

### A cuticular hair arising from the outside of the exoskeleton

### A cuticular process that is clearly articulated with the basal cuticle

### Setule

### A small bristle or spine on seta

### Sperm duct *(syn. Vas deferens)*

Narrow section of male reproductive system extending from posterior end of each testis to gonopore(s)

**Spinule**

Small spine along dorsal margin of carapace

**Tail spine** (*syn. apical spine, carapace spine, posterior spine, shell spine*)

Large, posteriorly directed, spine-like structure formed by posterodorsal extension of carapace

### Testis *(pl. testes)*

Paired section of male reproductive system in which sperm are produced, extend through anterior part of trunk, one to each side of midgut; opens to exterior via ventrally directed vas deferens

**Thoracic limb** (*syn. trunk limb, thoracic appendage, thoracopod*)

One of five pairs of appendages of anterior region of trunk (thorax); basically biramous, consisting of indistinct proximal protopodite (with endites and epipodite) and more distal endopodite and exopodite

**Thorax**

anterior limb-bearing region of trunk

**Trunk**

Posterior of two divisions of body (head, trunk), consist of thorax, abdomen, and postabdomen

**Transverse**

Crossing at right angles to the longitudinal axis; lying across or between

**Tubule (of maxillary gland)**

One of the convoluted ducts of paired maxillary gland

**Ventral**

The lower or underside of the body, that which is normally facing downwards; the side where the gap of the carapace is found

**References**

1. Crustacea Glossary by Natural History of Los Angeles County at <https://research.nhm.org/glossary/>
2. Maggenti, M. A. B. and Maggenti, A. R.; Scott L. Gardner (Ed) (2005): *Online Dictionary of Invertebrate Zoology.* <http://digitalcommons.unl.edu/onlinedictinvertzoology/2/> doi:10.13014/K2DR2SN5

**Table S2**: Tissue processing steps for serial dehydration and infiltration of *D. magna* samples with Formula R paraffin in tissue processor

| Duration | Solution | Temp (°C) | Vacuum (in. Hg) |
| --- | --- | --- | --- |
| 45 min | 80% Ethanol | 25 | 15 |
| 45 min | 95% Ethanol | 25 | 15 |
| 1 hour | 95% Ethanol | 25 | 15 |
| 1 hour (repeat thrice) | 100% Ethanol | 25 | 15 |
| 1 hour (repeat twice) | Xylene | 25 | 15 |
| 1 hour 30 min (repeat twice) | Paraffin | 60 | 15 |
| 2 hours | Paraffin | 60 | 15 |

**Table S3**: Automated steps for staining *D. magna* 5-μm sections with Harris’ hematoxylin and eosin in an auto-stainer

| Time | Solution |
| --- | --- |
| 3 min | Xylene |
| 5 min | Xylene |
| 2 min (repeat twice) | 100% Ethanol |
| 2 min | 95% Ethanol |
| 10 min | Tap water |
| 7 min | Hematoxylin |
| 1 min | Tap water |
| 1 min | Acidified Alcohol |
| 1 min | Tap water |
| 0.2 min | Ammoniated water |
| 1 min | Tap water |
| 0.3 min | Eosin |
| 1 min | 30% Ethanol |
| 1 min | 95% Ethanol |
| 1 min | 100% Ethanol |
| 1 min | Xylene |

**Table S4:** Anatomical ontology for DaHRA

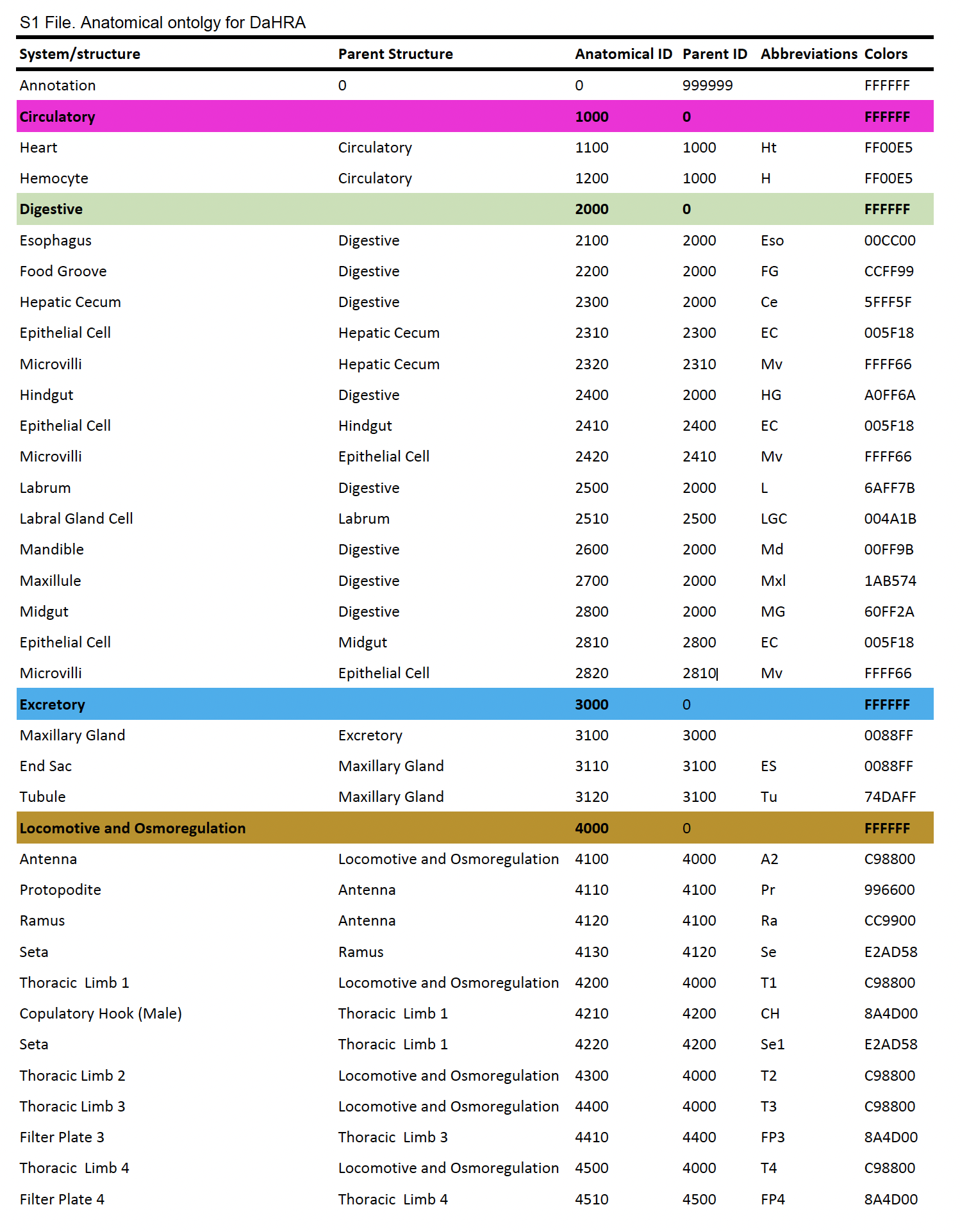

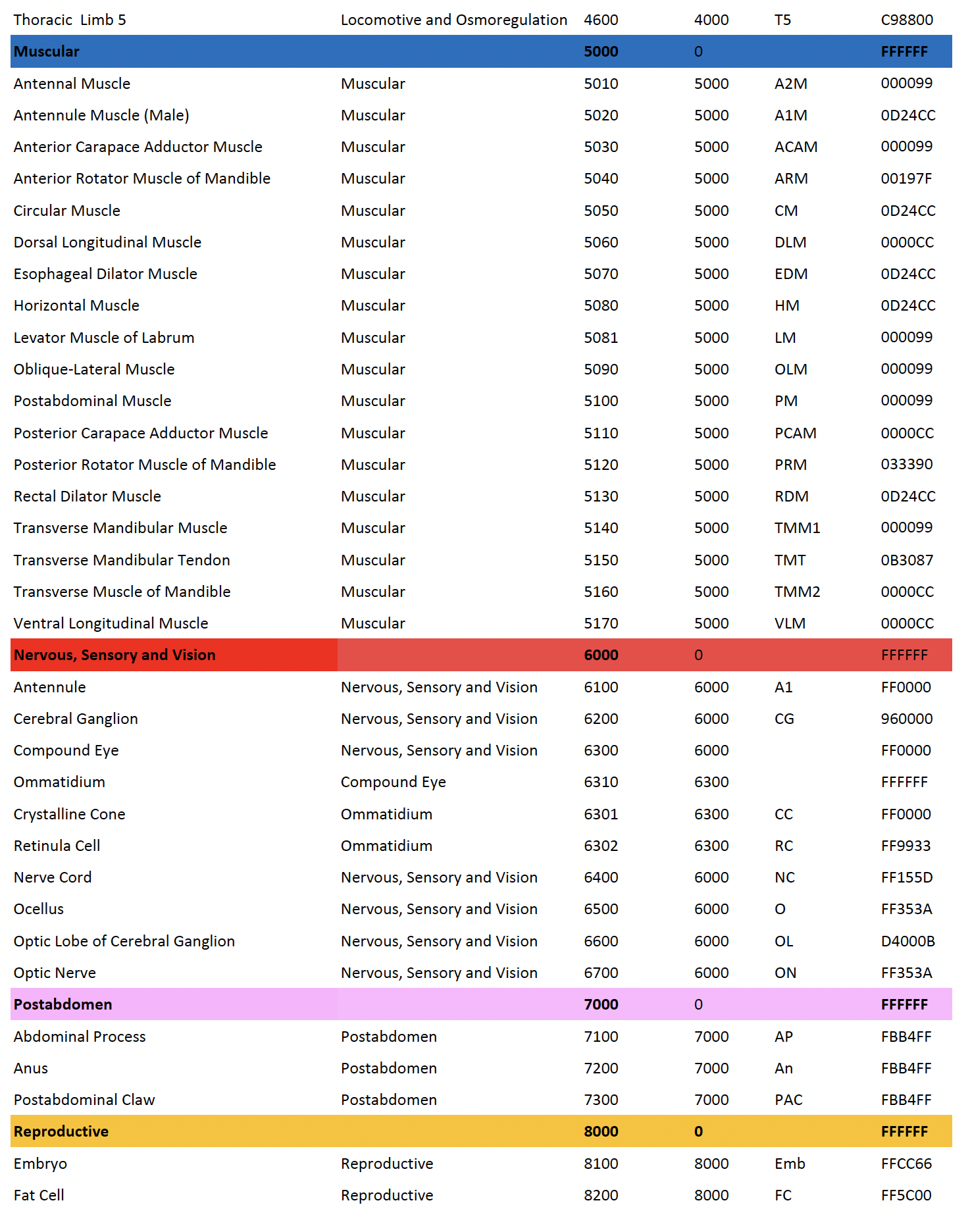

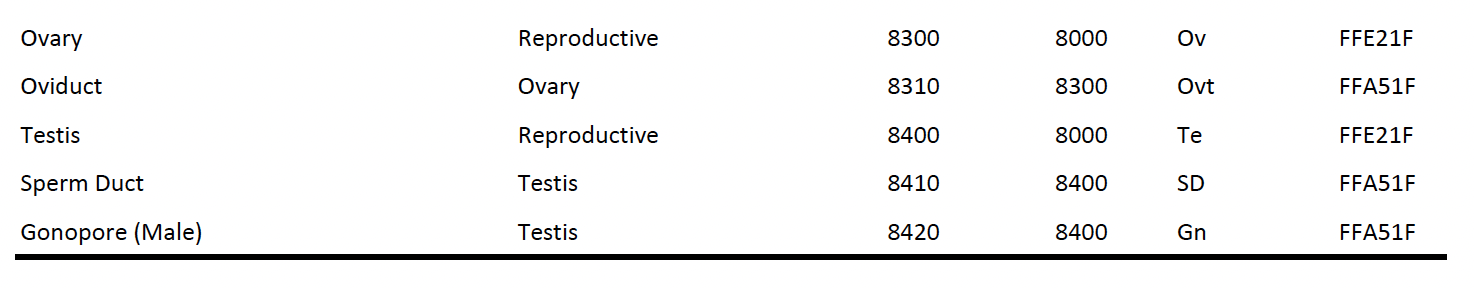

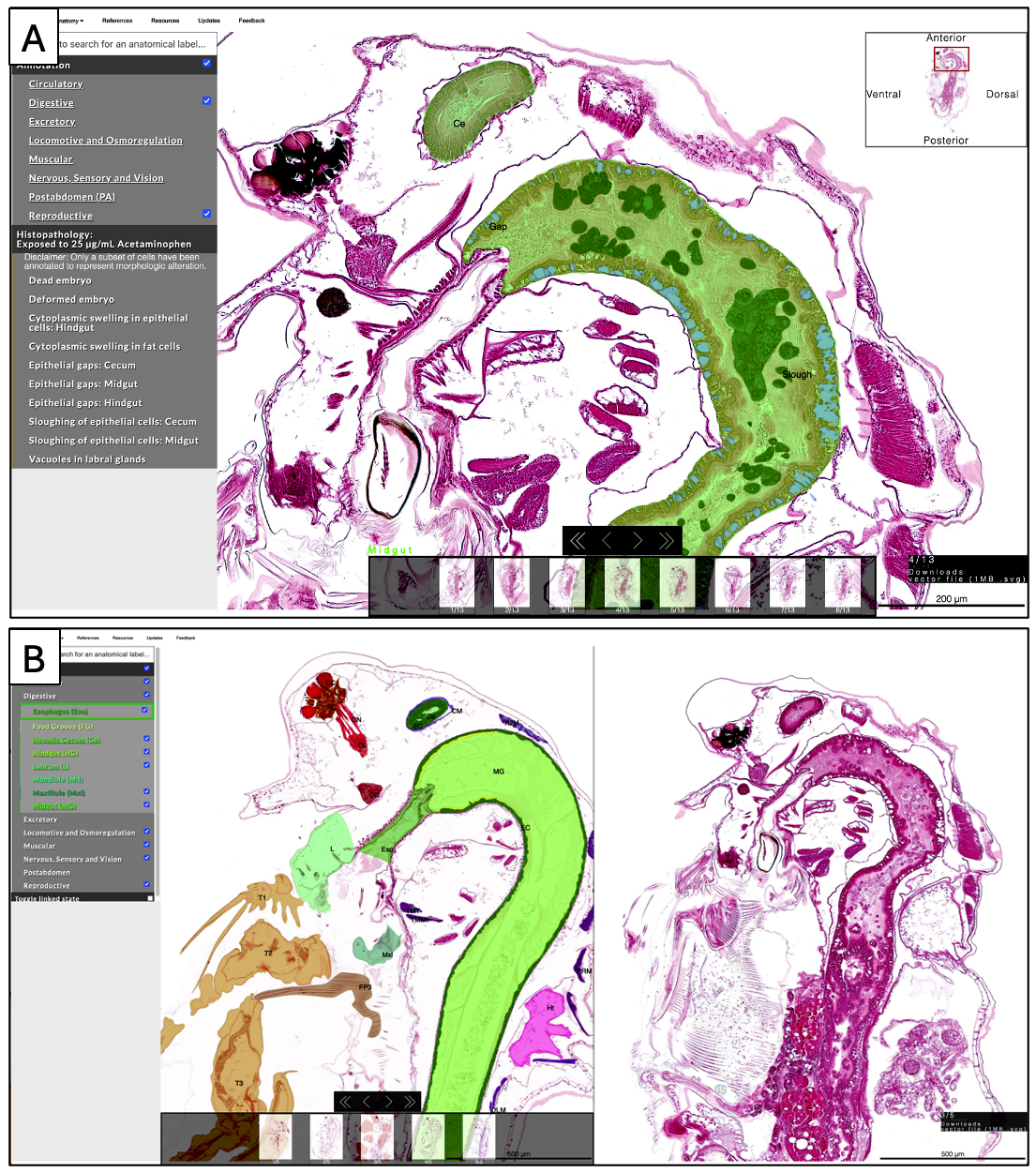

**Figure S5. Overview of DaHRA displaying annotated histopathological data. (A)** The affected organs or cell types in exposed *D. magna* are indicated by the checked boxes in the anatomical ontology and overlaid annotation in the images. The observed histopathologic features are listed under “Histopathology”. Clicking on each histopathologic feature will bring up the specific annotation. **(B)** Viewer comparing the wildtype (left) to the 25 µg/mL acetaminophen-exposed (right) *D. magna.*

**File S2**: Corresponding atlas links for each panel of Figure 2. Representative microanatomical structures of female and male *D. magna* in the three orthogonal planes.

A) <http://daphnia.io/anatomy/histology/?t=coronal_female&z=10>

B) <http://daphnia.io/anatomy/histology/?t=sagittal_female&z=10>

C) <http://daphnia.io/anatomy/histology/?t=transverse_female&z=10>

D) http://daphnia.io/anatomy/histology/?t=coronal_male&z=4

D’) <http://daphnia.io/anatomy/histology/?t=coronal_male&z=2&c=0.2,0.14,0.6,0.4>

E) <http://daphnia.io/anatomy/histology/?t=sagittal_male&z=3>

F) <http://daphnia.io/anatomy/histology/?t=transverse_male&z=8>

F’) <http://daphnia.io/anatomy/histology/?t=transverse_male&z=4&c=0.38,0.21,0.23,0.18>
